# A Two-Component System that Modulates Cyclic-di-GMP Metabolism Promotes *Legionella pneumophila* Differentiation and Viability in Low-Nutrient Conditions

**DOI:** 10.1101/604611

**Authors:** Elisa D. Hughes, Brenda G. Byrne, Michele S. Swanson

**Affiliations:** Department of Microbiology and Immunology, University of Michigan Medical School, Ann Arbor, Michigan, USA

**Author notes:** Address correspondence to Michele Swanson, PhD.

## Abstract

During its life cycle, the environmental pathogen *Legionella pneumophila* alternates between a replicative and a transmissive cell type when cultured in broth, macrophages, or amoebae. Within a protozoan host, *L. pneumophila* further differentiates into the hardy cell type known as the Mature Infectious Form (MIF). The second messenger cyclic-di-GMP coordinates lifestyle changes in many bacterial species, but its role in the *L. pneumophila* life cycle is less understood. Using an *in vitro* broth culture model that approximates the intracellular transition from the replicative to transmissive form, here we investigate the contribution to *L. pneumophila* differentiation of a two-component system (TCS) that regulates cyclic-di-GMP metabolism. The TCS is encoded by *lpg0278-lpg0277* and is co-transcribed with *lpg0279*, which encodes a protein upregulated in MIF cells. Using a *gfp*-reporter, we demonstrate that the promoter for this operon is RpoS-dependent and induced in nutrient-limiting conditions that do not support replication. The response regulator of the TCS (Lpg0277) is a bifunctional enzyme that both synthesizes and degrades cyclic-di-GMP. Using a panel of site-directed point mutants, we show that cyclic-di-GMP synthesis mediated by a conserved GGDEF domain promotes growth arrest of replicative *L. pneumophila*, production of pigment and poly-3-hydroxybutyrate storage granules, and viability in nutrient-limiting conditions. Genetic epistasis tests predict that the MIF protein Lpg0279 acts upstream of the TCS as a negative regulator. Thus, *L. pneumophila* is equipped with a regulatory network in which cyclic-di-GMP stimulates the switch from a replicative to a resilient state equipped to survive in low-nutrient environments.

**IMPORTANCE:** Although an intracellular pathogen, *L. pneumophila* has developed mechanisms to ensure long-term survival in low-nutrient aqueous conditions. Eradication of *L. pneumophila* from contaminated water supplies has proven challenging, as outbreaks have been traced to previously remediated systems. Understanding the genetic determinants that support *L. pneumophila* persistence in low-nutrient environments can inform design of remediation methods. Here we characterize a genetic locus that encodes a two-component signaling system (*lpg0278-lpg0277*) and a putative regulator protein (*lpg0279*) that modulates production of the messenger molecule cyclic-di-GMP. We show that this locus promotes both *L. pneumophila* cell differentiation and survival in nutrient-limiting conditions, thus advancing our understanding of the mechanisms that contribute to *L. pneumophila* environmental resilience.

## INTRODUCTION

*Legionella pneumophila* is a gram-negative bacterium commonly found in aquatic environments, where it replicates within protozoan hosts and persists within biofilms (1). When inhaled, contaminated water droplets transmit *L. pneumophila* to the human lung, where this opportunistic pathogen can infect alveolar macrophages. Studies examining the life cycle of *L. pneumophila* cultured in broth, macrophages, and amoebae support a developmental model in which nutrient levels govern cellular differentiation (2, 3). When nutrients are plentiful, the bacteria activate pathways that support growth; when nutrients become limiting, the progeny stop replicating and express multiple factors that promote *L. pneumophila* transmission to a new host, including flagella and the Dot/Icm Type IV secretion system (4). Within protozoan hosts, *L. pneumophila* can differentiate further to generate the resilient, metabolically dormant but highly infectious Mature Infectious Form (MIF), a cell-type believed to be prevalent in the environment (5, 6).

To alternate between replication within phagocytes and persistence within nutrient-poor aquatic environments, *L. pneumophila* relies on multiple regulatory mechanisms that coordinate rapid adaption to changing conditions (3). For example, replication in broth requires amino acids as the primary carbon source (7, 8), and a reduction in amino acid availability activates regulatory pathways that trigger conversion from the exponential (E) phase to the post-exponential (PE) transmissive phase (9, 10). The *L. pneumophila* life cycle is governed by a sophisticated regulatory network that includes the stringent response enzymes RelA and SpoT, multiple alternative sigma factors including RpoS and FliA, two-component regulatory systems, small regulatory RNAs, and CsrA post-transcriptional repressors (3, 11, 12). Driving the E to PE differentiation is a stringent response pathway coordinated by the alarmone guanosine penta- and tetraphosphate (abbreviated here as ppGpp) (13). The two-component system LetA/LetS responds to ppGpp accumulation by inducing transcription of small regulatory RNAs RsmY and RsmZ (14–16). These non-coding RNAs bind to and relieve repression by CsrA, enabling expression of multiple virulence traits associated with PE phase *L. pneumophila* including cytotoxicity, motility, and lysosome evasion (reviewed by 3).

Another second messenger molecule that regulates lifestyle switches in *L. pneumophila* and multiple other bacterial species is bis-(3’-5’)-cyclic dimeric guanosine (c-di-GMP) (4–8). Diguanylate cyclases (DCG), which possess a conserved GGDEF motif, synthesize c-di-GMP from two molecules of GTP. Conversely, phosphodiesterases (PDE) catalyze the hydrolysis of c-di-GMP back to GMP and contain either an EAL or HD-GYP domain (17). Most bacterial species utilizing c-di-GMP produce multiple enzymes that control c-di-GMP levels; for example, *Escherichia coli* encodes 29 proteins with GGDEF and/or EAL domains (18), and *Vibrio cholera* encodes over 60 such proteins (19).

The *L. pneumophila* genome (Philadelphia-1 and Lens strains) encodes 22 different enzymes involved in c-di-GMP metabolism, including several composite proteins possessing both GGDEF and EAL domains (20, 21). The range of activities influenced by these proteins is diverse and even includes control of opposing biological functions within the same cell. Recently Pecastings *et al*. (22) identified five c-di-GMP proteins in the Lens strain involved in biofilm regulation, three of which enhance biofilm formation while the other two inhibit this developmental program. In *L. pneumophila*, some c-di-GMP producing and degrading proteins enhance virulence by altering translocation of multiple Dot/Icm Type IV secretion system effectors, interfering with phagosome/lysosome fusion, and enhancing cytotoxicity—functions that promote replication within and transmission between host cells (20, 21). In general, GGDEF and/or EAL motifs are crucial for the protein’s enzymatic activity (21). However, genetic disruption of these domains does not always cause detectable changes in the cellular c-di-GMP concentration, leaving open the possibility that some of these proteins perform regulatory roles independently of c-di-GMP metabolism (20, 22).

Two-component regulatory systems, classically comprised of a histidine kinase and a response regulator, are a widespread signal transduction mechanism in bacteria that enables rapid adaptation to fluctuating conditions (reviewed by 23, 24). Some response regulators contain GGDEF and/or EAL domains; when phosphorylated on a conserved aspartate residue by its cognate histidine kinase, these enzymes can alter their c-di-GMP synthesis or hydrolysis (25). For example, in *Xanthomonas campestris*, the composite GGDEF/EAL protein RavR is the response regulator in a two-component system whose activation by the histidine kinase RavA shifts the enzyme from DCG to PDE activity and ultimately increases virulence (26). In the *L. pneumophila* Lens strain, Levet-Paulo et al. (27) characterized a putative two-component system comprised of a histidine kinase Lp0330 and its cognate response regulator Lpl0329, a bifunctional enzyme with both a GGDEF and an EAL domain. A series of *in vitro* experiments using purified proteins demonstrated that Lpl0329 possesses both DGC and the opposing PDE enzymatic activity, and phosphotransfer from Lpl0330 to Lpl0329 reduces DGC activity (27). In the *L. pneumophila* Philadelphila-1 strain, the homolog of Lpl0330 is Lpg0278 (hereafter HK), and the homolog of Lpl0329 is Lpg0277 (hereafter RR; HK and RR collectively are hereafter the TCS). On the Philadelphia-1 chromosome located directly 5’ of *lpg0278* and *lpg0277* is *lpg0279*, a gene encoding a hypothetical protein that is abundant in MIF cells (6).

The genetic proximity of the MIF gene *lpg0279* to the TCS-encoding *lpg0277* and *lpg0278* loci suggest potential co-regulation and related functions (28). As MIF cells are resilient forms that develop from PE phase *L. pneumophila* within protozoan hosts (10), here we test the hypothesis that the locus consisting of *lpg0279*, *lpg0278* and *lpg0277* (hereafter referred to as *lpg0279-77*) promotes persistence of *L. pneumophila* in nutrient-poor environments.

## RESULTS

### *lpg0277*, *lpg0278* and *lpg0279* are co-transcribed

The TCS-encoding genes *lpg0277* and *lpg0278* are located on the same DNA strand, 22 bp 3’ of *lpg0279*. To test the hypothesis that these three genes constitute an operon, we analyzed RNA extracted from wild-type (WT) *L. pneumophila* cultured to PE phase in rich AYET media. After conversion to cDNA, an endpoint PCR assay was conducted using primer sets designed to span the intergenic regions between the three genes (Fig. 1A). As predicted, amplicons of ∼300 bp were generated by primer sets A/B as well as by primers C/D, indicating that *lpg0279* and *lpg0278* form a single transcriptional unit, as do *lpg0278* and *lpg0277* (Fig. 1B). A similar experiment was conducted to determine if this transcriptional unit includes *lpg0280*, which is located 165 bp 5’ of *lpg0279* and codes for a putative transcriptional regulator of the LysR family. No product was generated for primer pairs E/F (Fig 1C), indicating that the operon consists solely of *lpg0279*, *lpg0278* and *lpg0277*.

**Fig. 1.**
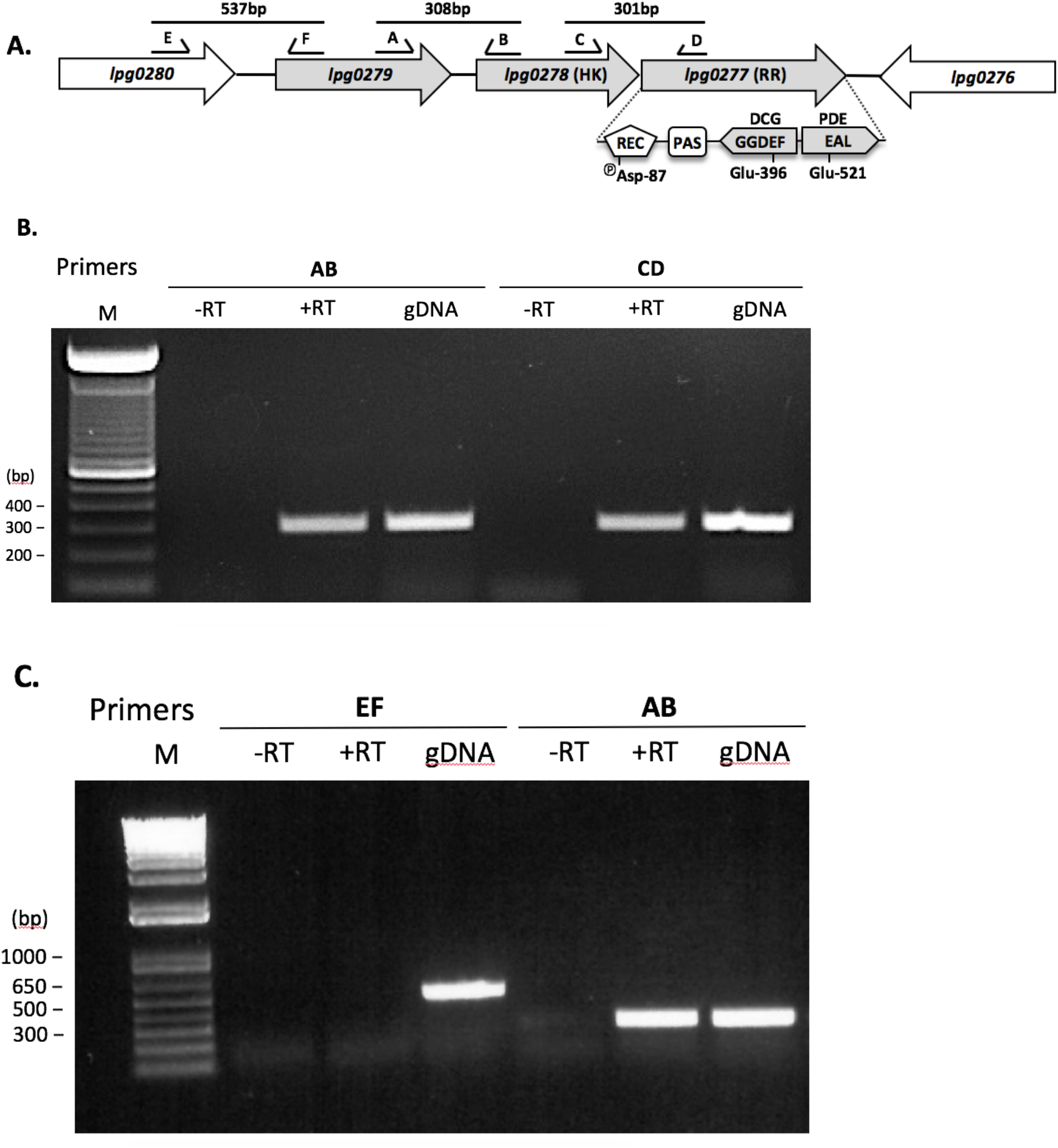
The genes *lpg0279*, *lpg0278* and *lpg0277* constitute an operon. **(A)** Schematic of the locus containing *lpg0279*, the TCS-encoding genes *lpg0278* and *lpg0277*, and the primer sets used to characterize mRNA by PCR. Also shown are the genes located 5’ and 3’ of the *lpg0279-2077* locus. Co-transcription of **(B)** *lpg0279, lpg0278* and *lpg0277* and **(C)** *lpg0280* and *lpg0279* and, as a positive control, *lpg0279* and *lpg0278* was assessed by end-point PCR assay with or without reverse transcriptase **(RT)** using RNA isolated from PE phase WT *L. pneumophila* that was converted to cDNA. As a reference for the PCR product length expected for each primer pair, genomic DNA **(gDNA)** was also used as a template. **M:** DNA size marker.

### The *lpg0279-77 locus* is induced at the transition from E to PE phase

To begin to delineate the role of *lpg0279-77* in the *L. pneumophila* life cycle, we first analyzed the timing and conditions that induce its promoter activity. To do so, we generated a transcriptional reporter by ligating a DNA fragment corresponding to the 832 bp immediately 5’ of the *lpg0279* open reading frame to a promoterless copy of the *gfp-mut3* gene encoded on plasmid pBH6119 (29), generating p*lpg0279-gfp*. This transcriptional reporter was then transferred to WT *L. pneumophila*.

Expression of *lpg0279-gfp* was monitored in E and PE phase broth cultures, which function as proxies for the intracellular replicative and transmissive stages, respectively (2, 4). In each case, E phase cultures were sub-cultured to an OD_600_ of 0.4-0.8 in rich AYE media, incubated at 37°C on an orbital shaker, and then GFP fluorescence and cell density were measured at 2-3 h intervals. As a reference for PE phase expression, a transcriptional reporter strain in which *gfp* expression is driven by the promoter for the flagellin subunit *flaA* (p*flaA-gfp*) was analyzed in parallel (13). Serving as the negative control was a strain carrying the promoterless *gfp* vector pBH6119. All strains grew equally well as measured by OD_600_, and no fluorescence was observed for the vector control (Fig. 2). Throughout E phase, the p*lpg0279-gfp* strain generated only minimal levels of GFP fluorescence, whereas promoter activity increased markedly upon entry into PE phase—kinetics similar to that of the p*flaA-gfp* marker of the PE transmissive phase (Fig. 2).

**Fig. 2.**
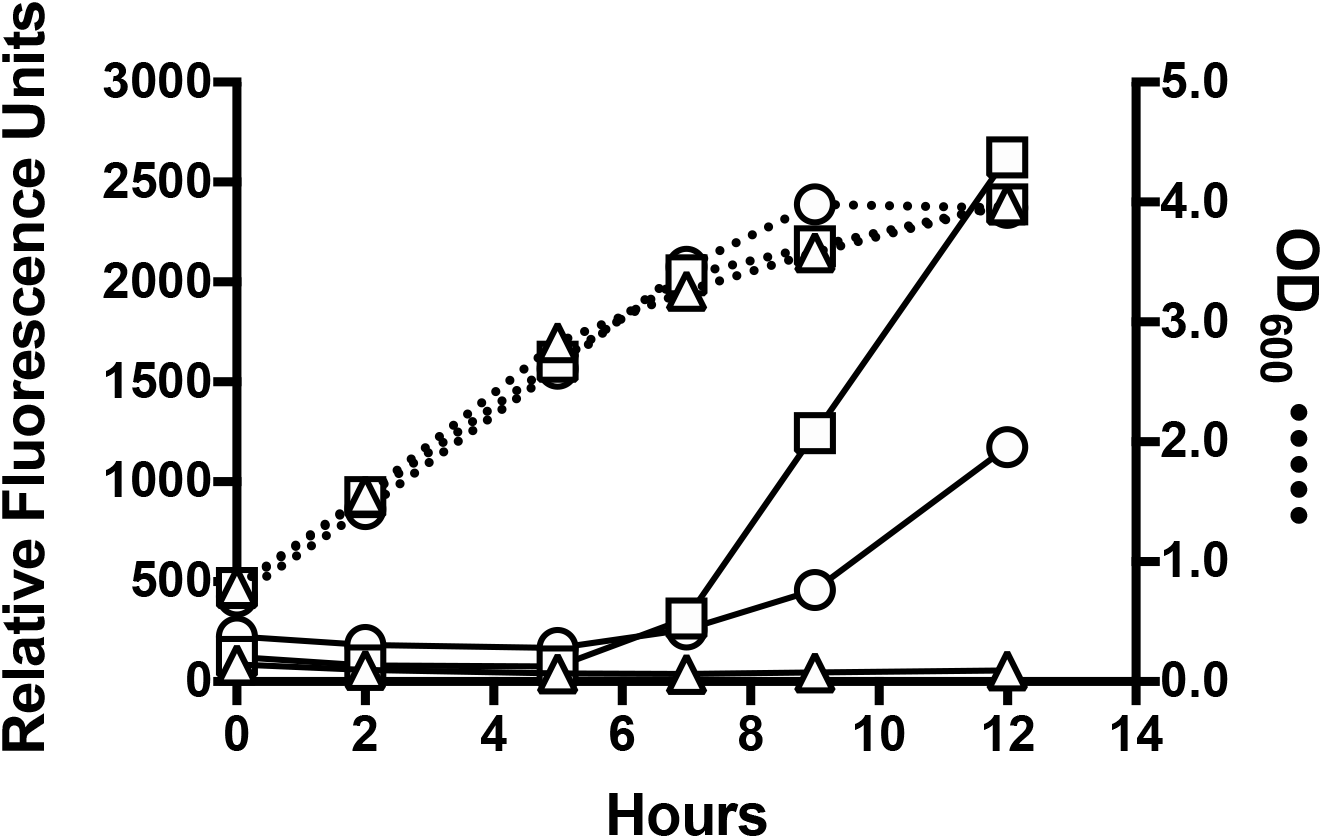
Promoter activity for the *lpg0279-77* operon increases upon entry into PE phase. GFP fluorescence generated by the transcriptional reporter p*lpg0279-gfp* (◯). Negative control strain is Lp02 carrying the empty pBH6119 vector (Δ); PE reference strain carries the flagellin subunit reporter p*flaA-gfp* (☐). For all strains, overnight E phase cultures were diluted to a starting OD_600_ of 0.8, incubated at 37°C, and then their fluorescence was measured at 2-3 h intervals. Relative Fluorescence Units (solid lines) was calculated as Fluorescence Units/OD_600_ (dotted lines) and represent the means ± SE of triplicate samples. In each case, error < 5%. Data shown are representative of results obtained in at least three independent experiments.

### The stationary phase sigma factor RpoS activates *lpg0279-77* expression

Due to the temporal similarity of their promoter activation (Fig. 2), we postulated that transcription of *lpg0279-77* and *flaA* may be controlled by the same regulatory proteins. Therefore, we assessed the contribution of a subset of the regulators known to coordinate the *L. pneumophila* transition from E to PE phase (16, 30–33). To do this, the p*lpg0279-gfp* and p*flaA-gfp* reporters were transformed into mutants lacking either the alternative sigma factors FliA (σ^28^) or RpoS (σ^S^/σ^38^), the two-component system LetA/S, or the ppGpp synthetase RelA. This panel of strains was then cultured on CYET agar at 37°C for 3 days. Visible differences in GFP expression indicated that RpoS is essential for robust transcription of *lpg0279-77*, but the other regulatory factors were not (Fig. 3A). In contrast, p*flaA-gfp* expression was only marginally diminished in the *rpoS* mutant; as expected, the FliA sigma factor was its critical regulator (Fig. 3B) (32, 34). Thus, although the promoters for *flaA* and *lpg0279-77* are each induced in PE phase, their mechanisms of regulation differ; accordingly, these two loci may respond to distinct environmental signals.

**Fig. 3.**
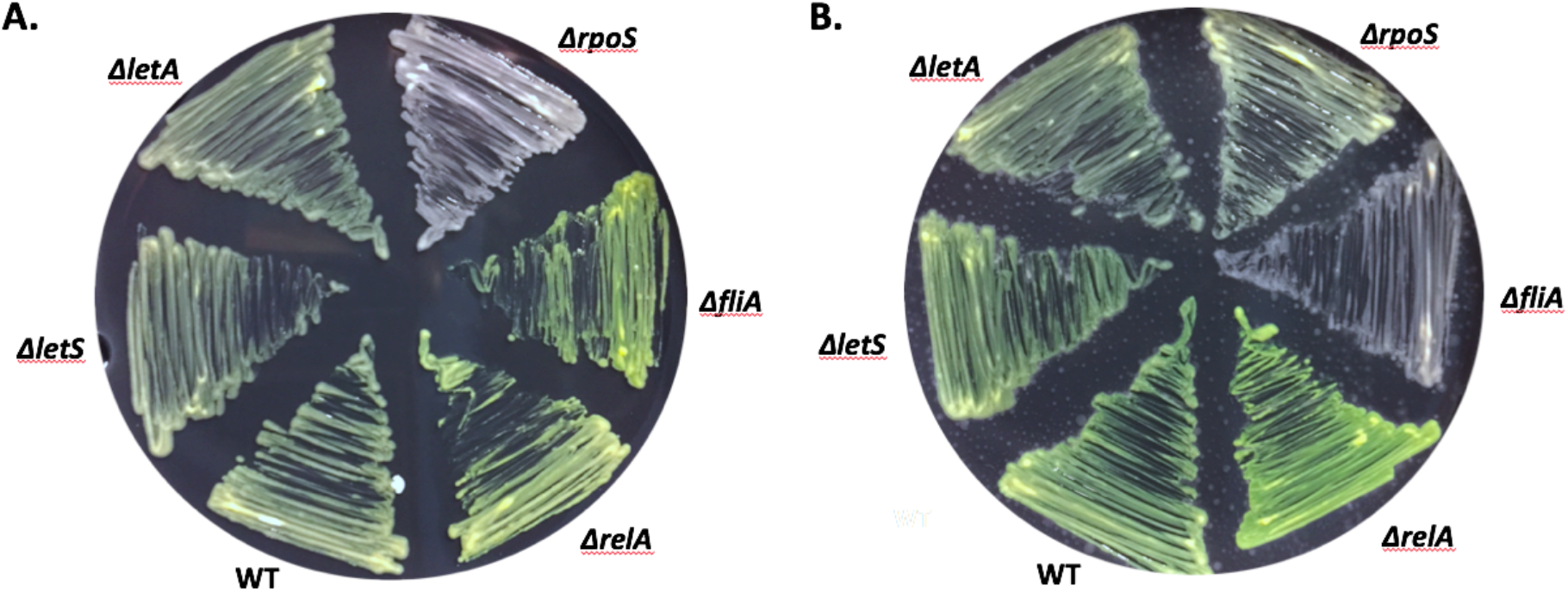
Expression of *lpg0279-77* is RpoS-dependent. Images obtained in ambient light of WT Lp02 and *letA*, *letS*, *relA*, *rpoS* and *fliA* mutants harboring the transcriptional reporter plasmids **(A)** p*lpg0279-gfp* or **(B)** p*flaA-gfp*, which serves as a reference for PE phase gene expression. The strains indicated were cultured on CYET at 37°C for 3 days to allow for bacterial growth and GFP accumulation.

### Expression of *lpg0279-77* increases in the absence of amino acids essential for replication

First identified in *E. coli* (35), the stationary phase sigma factor RpoS is widespread in proteobacteria (36). RpoS is a key regulator of the stringent response, which facilitates bacterial adaptation to a range of stresses, including starvation (37). When nutrients become limiting, replicating *L. pneumophila* accumulate the alarmone ppGpp and synthesize RpoS, which activates expression of multiple genes critical for fitness in the PE phase (3).

Because *lpg0279-77* transcription is RpoS-dependent, we next examined whether *lpg0279-gfp* expression by replicating bacteria was induced when nutrients were limited. Standard growth media for *L. pneumophila* consists of a rich media (AYE) supplemented with both iron and the amino acid L-cysteine, as this bacterium lacks a number of cysteine biosynthesis enzymes (38, 39). Accordingly, we first quantified *lpg0279-gfp* fluorescence in *L. pneumophila* cultured in AYE containing both L-cysteine and ferric nitrate, either L-cysteine or ferric nitrate alone, or neither supplement. In media supplemented with L-cysteine, either with or without additional iron, *L. pneumophila* continued to replicate for > 9 hours and did not activate the *lpg0279-77* promoter (Fig. 4A). In contrast, when cultures lacked L-cysteine, bacterial replication stalled and the *lpg0279-77* promoter was induced (Fig. 4A).

**Fig. 4.**
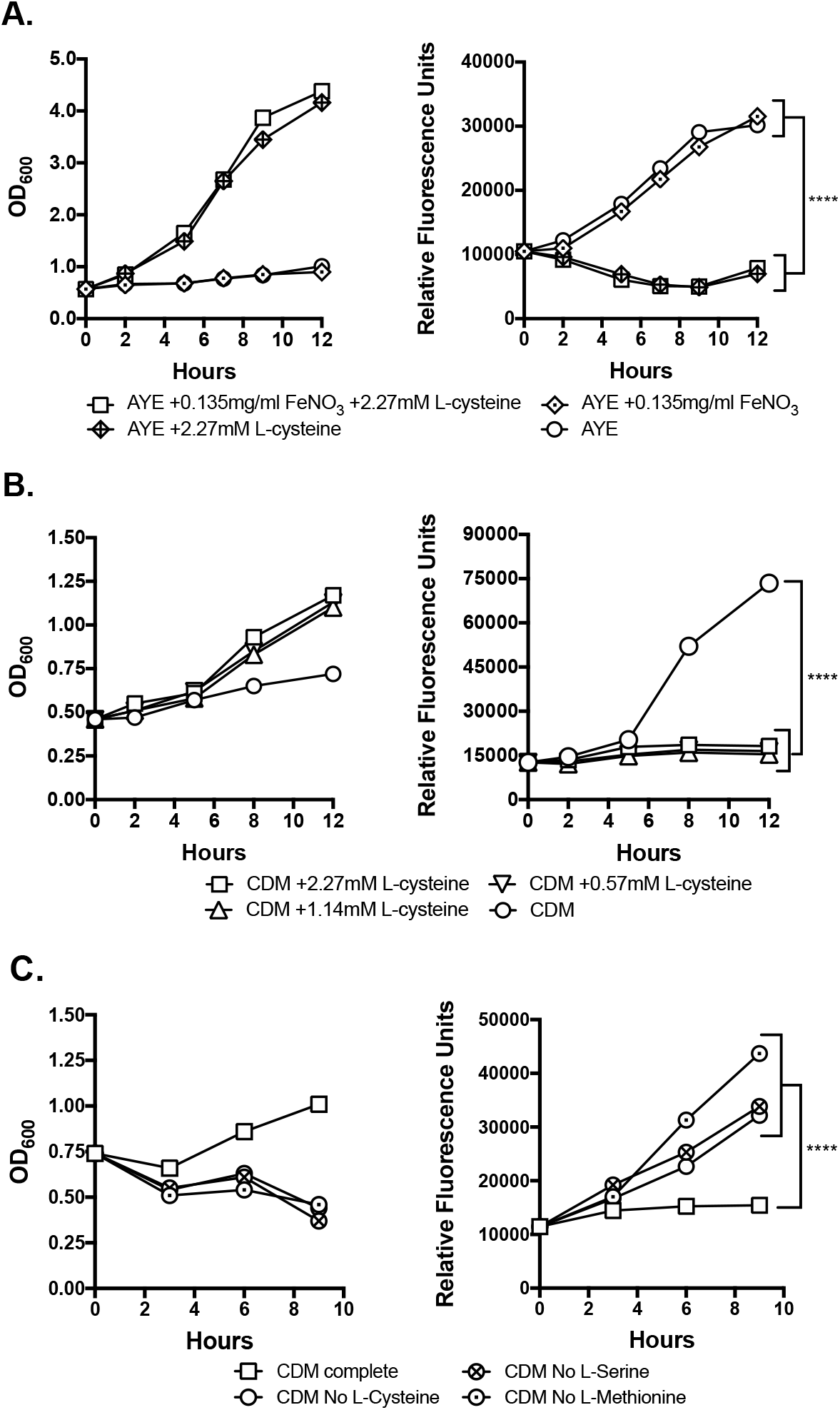
Nutrient limitation induces expression of *lpg0279-77*. To examine the impact of nutrient limitation on *lpg0279-77* transcription, E phase Lp02 cultures carrying the p*lpg0279-gfp* reporter plasmid were exposed to culture conditions shown and incubated for 9-12 h at 37°C on an orbital shaker, with GFP fluorescence and cell density (OD_600_) measured at 2-3 h intervals. **(A)** OD_600_ measurements and RFU values obtained for cultures exposed to AYE medium with or without 0.135 mg/ml ferric nitrate and/or 2.27 mM L-cysteine. **(B)** OD_600_ measurements and RFU values obtained for cultures exposed to CDM without or with the indicated concentration of L-cysteine. **(C)** OD_600_ and RFU values obtained for cultures exposed to CDM with or without either L-cysteine, L-methione, or L-serine. RFU symbols represent the means ± SE of triplicate GFP fluorescence readings, normalized to OD_600_ values obtained by measuring a 1/10 dilution of cell culture in a spectrophotomer (short error bars are masked by symbols). A two-tailed Student’s *t*-test was used to determine statistically significant differences between groups at 12 h (****, *p* < 0.0001). Data shown are representative of results obtained in two or more independent experiments.

The yeast extract in AYE contains several amino acids including L-cysteine (bdbiosciences.com), so to more accurately address the impact of this amino acid in promoting *lpg0279-77* expression we repeated the analysis using a chemically defined medium (CDM) (38) in which L-cysteine, L-cystine, and supplemental ferric pyrophosphate were omitted. Initial experiments examined *lpg0279-gfp* fluorescence in CDM containing a range of L-cysteine concentrations: 100% (2.27 mM), 50% (1.14 mM) and 25% (0.57 mM) of the standard concentration used to support *in vitro* growth in rich AYE media. In each case, the presence of L-cysteine repressed expression of *lpg0279-gfp* by replicating *L. pneumophila* (Fig. 4B).

These experiments also revealed an inverse relationship between *lpg0279-77* promoter activity and bacterial growth (Figs. 4A-B; also see Fig. 2). Specifically, the absence of L-cysteine hindered bacterial replication and induced *lpg0279-77* gene expression. One interpretation is that *lpg0279-77* transcription is triggered by the absence of this particular amino acid; in turn, the locus suppresses replication. Alternatively, the inability of *L. pneumophila* to replicate in the absence of an essential amino acid—in this case L-cysteine—may be a signal that induces *lpg0279-77* expression. To distinguish between these two possibilities, we examined *lpg0279-gfp* fluorescence in CDM lacking either L-serine or L-methionine, two amino acids *L. pneumophila* requires for growth in CDM (38). Compared to cultures in complete CDM, lack of either L-serine or L-methionine reduced replication and enhanced p*lpg0279-gfp* expression (Fig. 4C). Thus, *L. pneumophila* induces *lpg0279-77* promoter activity in response to nutrient-limiting conditions that impede bacterial replication.

### PE phase *L. pneumophila* lacking either the HK or RR component of the TCS, or constitutively expressing *lpg0279*, exhibit a shortened lag phase and reduced pigmentation

Based on the increase in *lpg0279-77* promoter activity observed in response to conditions that do not support *L. pneumophila* growth (Figs. 2 and 4), we next examined whether this locus influences differentiation of replicating *L. pneumophila* to the PE phase. To do so, we generated isogenic mutants containing in-frame deletions in either *lpg0279*, *lpg0278*, or *lpg0277*.

To assess growth of each mutant strain, overnight E phase (OD_600_ < 2.5) or PE phase (OD_600_ < 3.5) cultures were diluted to an OD_600_ of 0.1 in AYET, and then cell density was quantified over a 36 h period using a Bioscreen growth curve analyzer. For the E phase inocula, growth curves for each mutant resembled the WT strain (data not shown). However, for the PE phase inocula, mutants lacking either the HK (Δ*lpg0278*) or RR (Δ*lpg0277*) component of the TCS mimicked the WT E phase reference culture by exhibiting a minimal lag phase (Fig. 5A and B). Although expression of each individual WT gene from an IPTG-inducible promoter was insufficient to remedy this defect (data not shown), IPTG-induced expression of the *lpg0278-lpg0277* locus from plasmid pHK/RR restored WT growth kinetics (Fig. 5A and B).

**Fig. 5.**
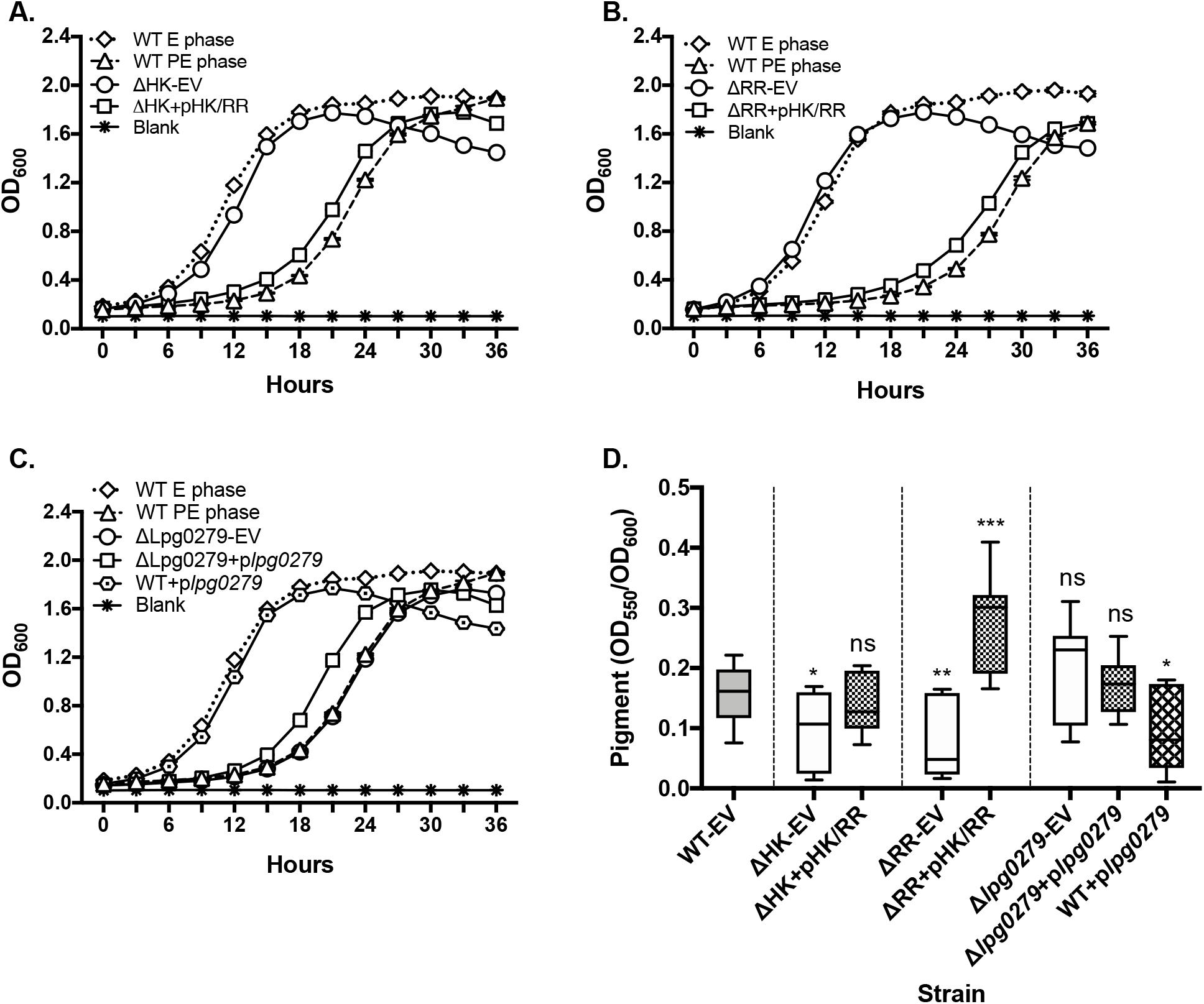
*L. pneumophila* lacking a complete TCS or ectopically expressing *lpg0279* resemble WT E phase cells. The growth kinetics of WT *L. pneumophila* inocula in E phase (dotted lines) or PE phase (dashed lines) was compared to PE phase inocula of **(A)** an ΔHK mutant and its complement, **(B)** an ΔRR mutant and its complement, and **(C)** an Δ*lpg0279* mutant and its complement, together with a WT strain of *L. pneumophila* constitutively expressing *lpg0279*. With the exception of the WT E phase reference culture (OD_600_ < 2.0), all strains were cultured overnight in AYET medium to OD_600_ > 3.5 (corresponding to WT PE phase cultures), then diluted to a starting OD_600_ of ∼0.1 and incubated for 36 h in a Bioscreen growth curve analyzer set at 37°C with continuous shaking; OD_600_ measurements were obtained at 3 h intervals. Shown are means ± SE of triplicate samples, and data shown are representative of three independent experiments. (**D**) Pigment accumulation in late PE phase cultures of ΔHK, ΔRR and Δ*lpg0279* mutants and the corresponding complemented strains, and of WT L. *pneumophila* constitutively expressing *lpg0279*. Supernatants of broth cultures maintained in PE phase for 1 or 3 days were collected by centrifugation, their absorbance at OD_550_ quantified and then normalized to cell density (OD_600_). Results shown are the means ± SE of pooled data from three independent experiments, with duplicate readings obtained for each measurement. A two-tailed Student’s *t*-test was used to determine statistically significant differences in pigmentation compared to WT-EV (**ns**: not significant; **\***, *p* <0.05; **\*\***, *p* < 0.01; **\*\*\***, *p* < 0.001). **EV:** strain harbors the pMMB206cam empty vector.

In contrast to the TCS genes, growth of mutants lacking *lpg0279* was indistinguishable from WT PE phase cultures (Fig. 5C). However, WT *L. pneumophila* constitutively expressing a plasmid-borne allele of *lpg0279* engineered to encode an optimal ribosome binding site (20) exhibited growth kinetics similar to the ΔHK or ΔRR mutants and E phase WT *L. pneumophila* (Fig. 5C). Therefore, constitutive expression of *lpg0279* inhibits replicating *L. pneumophila* from transitioning to PE phase, whereas the genetically-linked TCS promotes differentiation of replicating *L. pneumophila* to the PE phase.

To test more rigorously the impact of the TCS and Lpg0279 on *L. pneumophila* differentiation, we quantified production of the soluble pigment pyomelanin, a late PE phase trait (40, 41). Derived from secreted homogentisic acid (HGA), this melanin-like substance is not required for intracellular survival; rather it enhances environmental fitness of *L. pneumophila* by protecting bacterial cells from the damaging effects of light and by aiding in iron acquisition through its ferric reductase activity (42, 43). When cultured in rich broth to a cell density typical of PE phase (OD_600_ > 3.5), all strains generated minimal pigment. However, when maintained in PE phase for up to three days, WT cultures accumulated pigment, but strains that lacked either of the TCS components did not (Fig. 5D). Consistent with their growth phenotypes, deletion of the *lpg0279* gene had no effect, whereas constitutive expression of *lpg0279* by WT *L. pneumophila* inhibited pigment accumulation (Fig. 5D). Consistently, the ΔRR mutant harboring the complementing plasmid pHK/RR produced higher levels of pigment than did the WT strain, indicating that the TCS in multi-copy may stimulate differentiation to PE phase. As TCS integrity appears essential for the transition from E to PE phase in *L. pneumophila*, we next examined whether the TCS enhances *L. pneumophila* viability when nutrients are limiting. In TCS signal transduction pathways, RR activity is distal to HK; accordingly, we analyzed the RR mutant as representative of Lpg0278-0277 TCS output.

### The TCS facilitates PHB production and long-term survival in low-nutrient conditions

Beginning in the PE phase, *L. pneumophila* generates large poly-3-hydroxybutyrate (PHB) inclusions, a reserve carbon and energy source that accumulates in MIF cells (5, 44) and enhances persistence of *L. pneumophila* in low-nutrient environments (45). Therefore, we next quantified the PHB content of WT and the ΔRR mutant using the lipophilic dye Nile Red, a fluorescent stain that is highly specific for intracellular lipids, including PHB (46). To determine the baseline value, the fluorescence of E phase cultures in AYET was quantified. Next, after collecting WT and ΔRR mutant cells by centrifugation, their expression of *lpg0279-77* was induced by resuspending each cell sample in CDM media lacking L-cysteine (Fig. 4B), and then the bacteria were incubated for 24 h at 37°C with aeration before a second PHB quantification. As expected, in E phase the Nile Red PHB signal for both WT and ΔRR was negligible. However, after 24 h of nutrient limitation, the WT cells exhibited significantly greater fluorescence than did the ΔRR mutant (Fig. 6A). Expression by the mutant of the TCS from pHK/RR not only fully remedied this defect, but also generated a PHB signal exceeding that of the WT strain. Therefore, the TCS promotes accumulation of PHB storage granules in *L. pneumophila*.

**Fig. 6.**
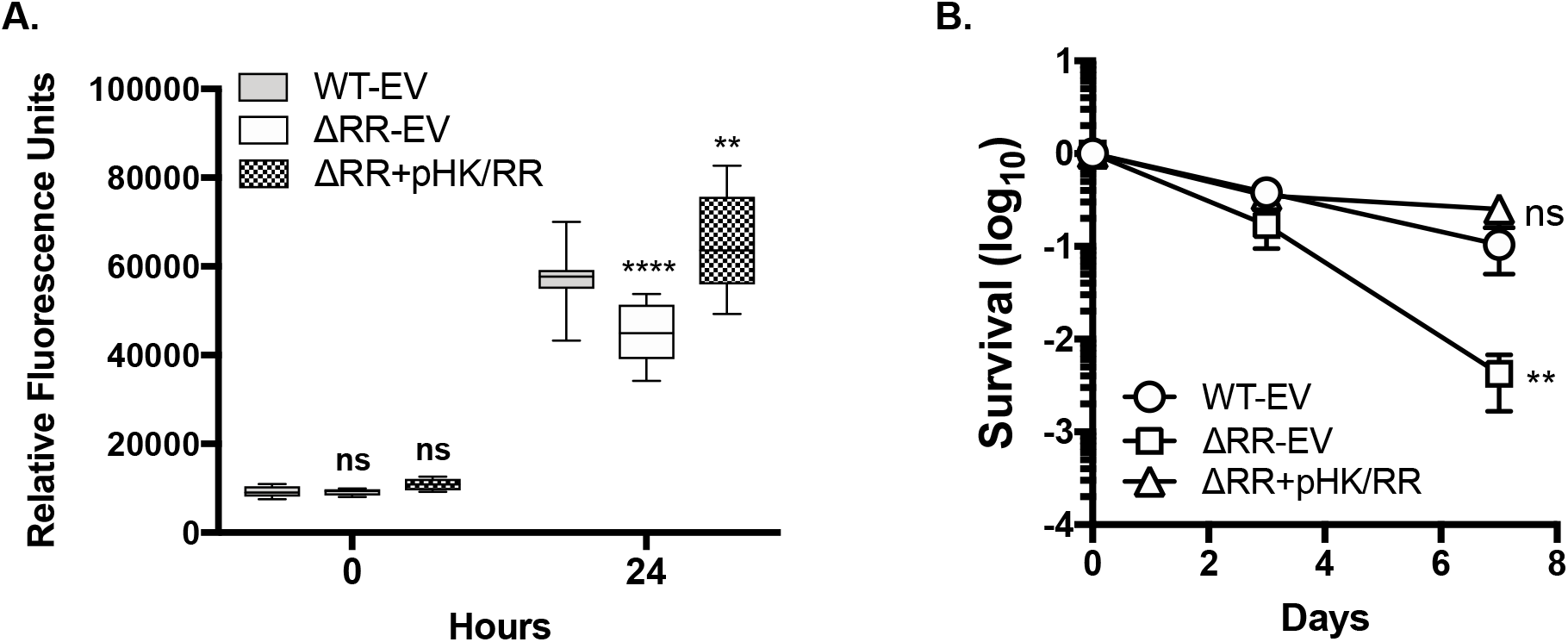
The TCS promotes PHB production and long-term viability. E phase cultures of WT *L. pneumophila*, the ΔRR mutant, and the ΔRR complemented strain harboring pHK/RR were collected by centrifugation, resuspended in CDM lacking L-cysteine, and incubated at 37°C on an orbital shaker for up to 7 d. **(A)** Quantification of PHB content before and after 24 h CDM exposure. Results shown are the means ± SE of pooled data obtained from triplicate samples in four independent experiments. A two-tailed Student’s *t*-test was used to determine statistically significant differences in fluorescence compared to WT-EV (**ns**: no significance; **\*\***, *p* < 0.01; **\*\*\*\***, *p* < 0.0001). **(B)** Survival was quantified by plating serial dilutions of the cultures indicated and enumerating CFUs before (titer) and after 3 and 7 d incubation. Shown are ratio of CFU(day)/CFU(titer), with symbols representing the means ± SE of pooled data obtained from duplicate samples in four independent experiments. The Mann-Whitney test was used to determine statistically significant differences in survival compared to WT-EV (**ns**: no significance; **\*\***, *p* < 0.01). **EV:** strain harbors the pMMB206cam empty vector.

As this TCS equips replicating *L. pneumophila* to respond to nutrient limitation by transitioning to the PE phase, producing pigment, and accumulating PHB storage granules—all traits that increase resilience in the environment—we next investigated whether this TCS facilitates *L. pneumophila* survival during prolonged exposure to low-nutrient conditions. To do so, we quantified CFUs of WT and ΔRR mutant cells first in E phase and then again after 3 and 7 d incubation in CDM lacking L-cysteine, as described for the Nile Red fluorescence experiments. Indeed, compared to WT, the ΔRR strain suffered a greater loss of viability by day 7, a defect remedied by ectopic expression of the TCS locus (Fig. 6B). Therefore, *L. pneumophila* persistence in nutrient-limited conditions is enhanced by the *lpg0277* locus, which encodes an enzyme equipped to modulate levels of the second messenger c-di-GMP.

### The GGDEF domain of Lpg0277 promotes transition from E to PE phase

Because the RR is a bifunctional enzyme with both DGC and PDE activity (27), we next examined which of these opposing functions accounts for the ΔRR mutant phenotypes. To do so, we took advantage of the known contributions of the GGDEF and EAL amino acid motifs to DCG and PDE activity, respectively (17, 25). To abrogate DGC function, we engineered point mutant strains in which the conserved Glu-396 residue was replaced with Lys (27), generating the RR^E396K^ allele. Likewise, to impair PDE activity, the Glu-521 residue in the EAL domain was replaced with Ala, creating the RR^E521A^ allele (18). After confirming the DNA sequence of each *L. pneumophila* chromosomal point mutation, the corresponding mutant strains were transformed with either the complementing plasmid pHK/RR or the empty vector.

We first examined the growth kinetics and pigment production of the RR^E396K^ DCG and RR^E521A^ PDE point mutants after culturing to an OD_600_ > 3.5, correlating with PE phase in WT cells. The RR^E521A^ PDE mutant mimicked PE phase WT cells in growth kinetics (Fig. 7B), and its pigment production exceeded that of the WT strain (Fig. 7C). In contrast, the RR^E396K^ DCG mutant resembled E phase WT cells, as judged by its minimal lag phase (Fig. 7A) and decreased pigmentation (Fig. 7C), two defects that were complemented by ectopic expression of the WT TCS. Thus, the GGDEF motif of the RR promotes the transition of replicating *L. pneumophila* to the PE phase.

**Fig. 7.**
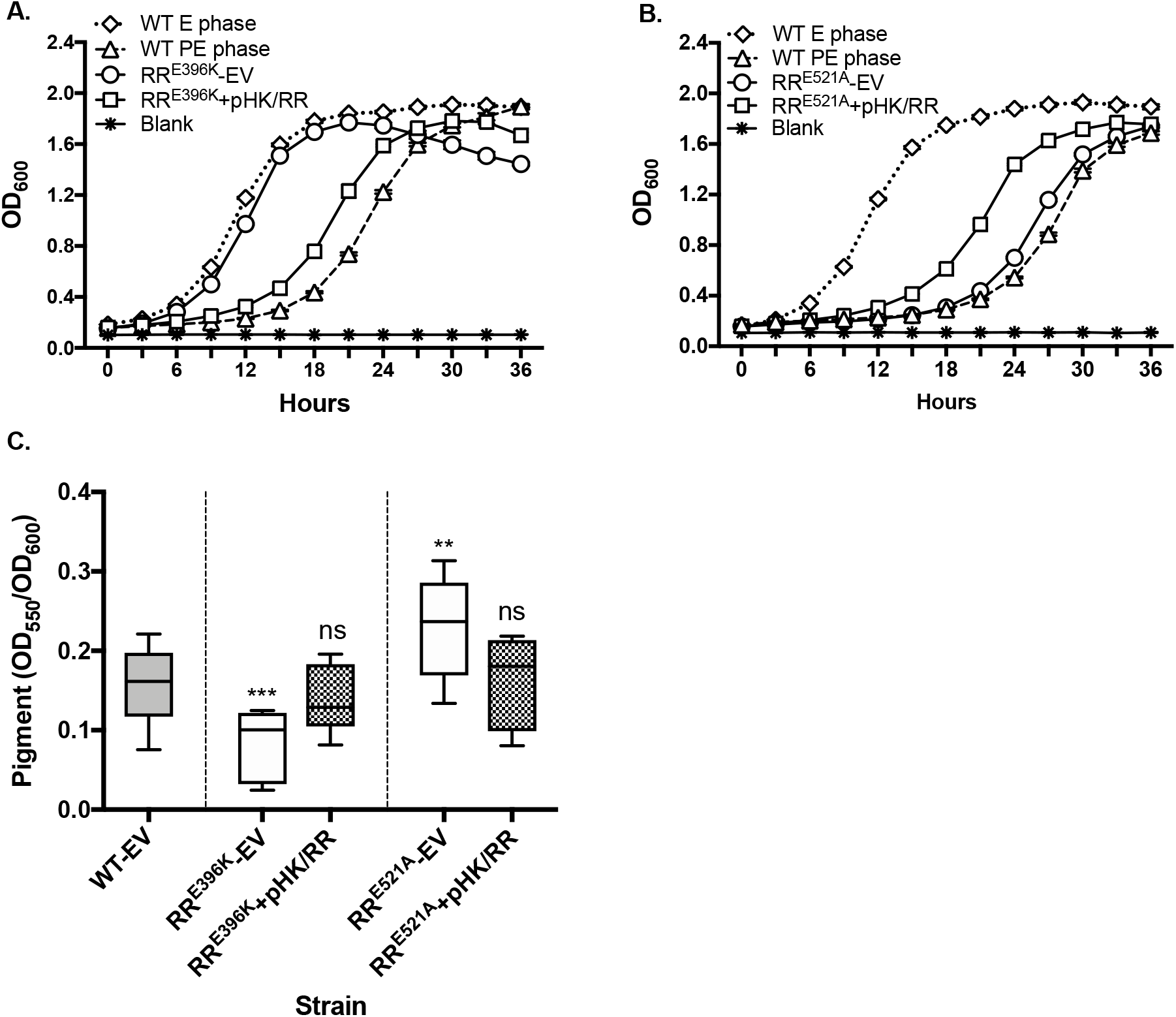
The DCG activity of RR Lpg0277 promotes transition to PE phase. Growth kinetics in AYET of WT *L. pneumophila* inocula in E (dotted lines) or PE phase (dashed lines) was compared to PE phase inocula of **(A)** RR^E396K^ point mutant and **(B)** RR^E521A^ point mutant strains. Symbols denote the means ± SE of triplicate samples (short error bars are masked by symbols), and data are representative of three independent experiments. **(C)** Pigment accumulation by WT *L. pneumophila* after maintenance in PE phase for 1-3 days, compared with the RR^E396K^ DGC and RR^E521A^ PDE point mutants and their respective complemented strains. Results shown are the means ± SE of pooled data obtained from duplicate samples in three independent experiments. A two-tailed Student’s *t*-test was used to determine statistically significant differences in pigmentation compared to WT (**ns**: not significant; **\*\****p* < 0.01; **\*\*\***, *p* < 0.001). **EV:** strain harbors the pMMB206cam empty vector.

To further probe RR function, we genetically abrogated the ability of the RR to be phosphorylated by its cognate HK, a post-translational modification that induces a change in RR enzymatic activity (27). For this purpose, the conserved Asp-87 residue in the putative phosphoacceptor site of the RR was replaced with Asn, generating mutant strain RR^D87N^ (27). The RR^D87N^ phosphoacceptor mutant exhibited growth and pigmentation defects similar to that observed for both the ΔRR and RR^E396K^ DGC mutants (Fig. S1; also Figs. 5 & 7), indicating a functional link between TCS phosphorylation and RR DGC activity.

Based on our growth and pigmentation analyses, we investigated whether abrogated DGC activity accounted for the reduction in both PHB levels and viability of ΔRR mutant cells (Fig. 6). To test this hypothesis, we exposed E phase cultures of the RR^E396K^ DGC, RR^E521A^ PDE, and RR^D87N^ phosphoacceptor point mutants to CDM lacking L-cysteine, and then quantified PHB production via Nile Red florescence and survival via CFU enumeration, as described for the ΔRR mutant analysis (Fig. 6). As expected, PHB accumulation by the RR^E521A^ PDE mutant was indistinguishable from WT *L. pneumophila*, and both strains survived well after 7 d exposure to CDM lacking L-cysteine (Fig. S2A). However, the RR^E396K^ DCG and RR^D87N^ phosphoacceptor point mutants each had reduced PHB content, as judged by Nile Red fluorescence (Figs. S2B-C). Both strains also lost viability after extended CDM exposure, a phenotype mimicking the ΔRR strain (Fig. S2B-C and 6B). It is notable that these defects were only partially complemented, perhaps due to unknown effects of perturbed cellular c-di-GMP pools. Nevertheless, the phenotypic profile of each point mutant is consistent with a model in which DCG activity and concomitant accumulation of cyclic-di-GMP stimulates replicating *L. pneumophila* to transition to PE phase and generate PHB stores that support bacterial survival in nutrient-limited conditions.

### Ectopic expression of *lpg0279* counteracts TCS function

The spatial proximity and co-regulation of *lpg0279* with the TCS-encoding genes *lpg0277* and *lpg0278* (Fig. 1) suggest a regulatory interaction. Since WT cells constitutively expressing *lpg0279* phenocopy the growth and pigmentation defects of the ΔRR and RR^E396K^ DCG mutants (Figs. 5 & 7), we hypothesized that Lpg0279 functions as a negative regulator of TCS activity.

To investigate whether Lpg0279 acts upstream of the TCS, the RR^E521A^ PDE point mutant—which resembles WT in the transition from E to PE phase (Fig. 7B-C)—was transformed with plasmid p*lpg0279* and then treated with IPTG to induce constitutive expression. In parallel, we also transformed the RR^E396K^ mutant with p*lpg0279* to evaluate any additive effects of loss of DGC activity and gain of Lpg0279 function. After culturing both strains to an OD_600_ > 3.5, we performed growth curve and pigmentation analyses. As expected, expression of *lpg0279* did not rescue the growth defect of the RR^E396K^ DCG mutant strain (Fig. S3A). However, *lpg0279* expression significantly shortened the lag phase of the RR^E521A^ PDE mutant to that of WT E phase cells (Fig. S3B). Assessment of pigment production yielded similar results, with *lpg0279* expression reducing pigment levels in the RR^E521A^ mutant but having no effect on the RR^E396K^ mutant cells (Fig. 8A). Furthermore, when exposed to CDM lacking L-cysteine for 7 d, both the WT and RR^E521A^ PDE mutant strains harboring p*lpg0279* suffered a significant drop in cell viability compared to the respective parent strains (Fig. 8B). These data indicate that *lpg0279* is epistatic to the TCS genes, acting as a negative regulator. In low-nutrient conditions, and in the presence or absence of an inducing signal, repression by Lpg0279 is relieved, enhancing DCG activity of the RR and increasing c-di-GMP. Although the downstream effectors of c-di-GMP generated by the RR remain to be identified, the activity of this TCS stimulate *L. pneumophila* to switch from a replicative state to a more resilient cell type better equipped to survive in low-nutrient environments.

**Fig. 8.**
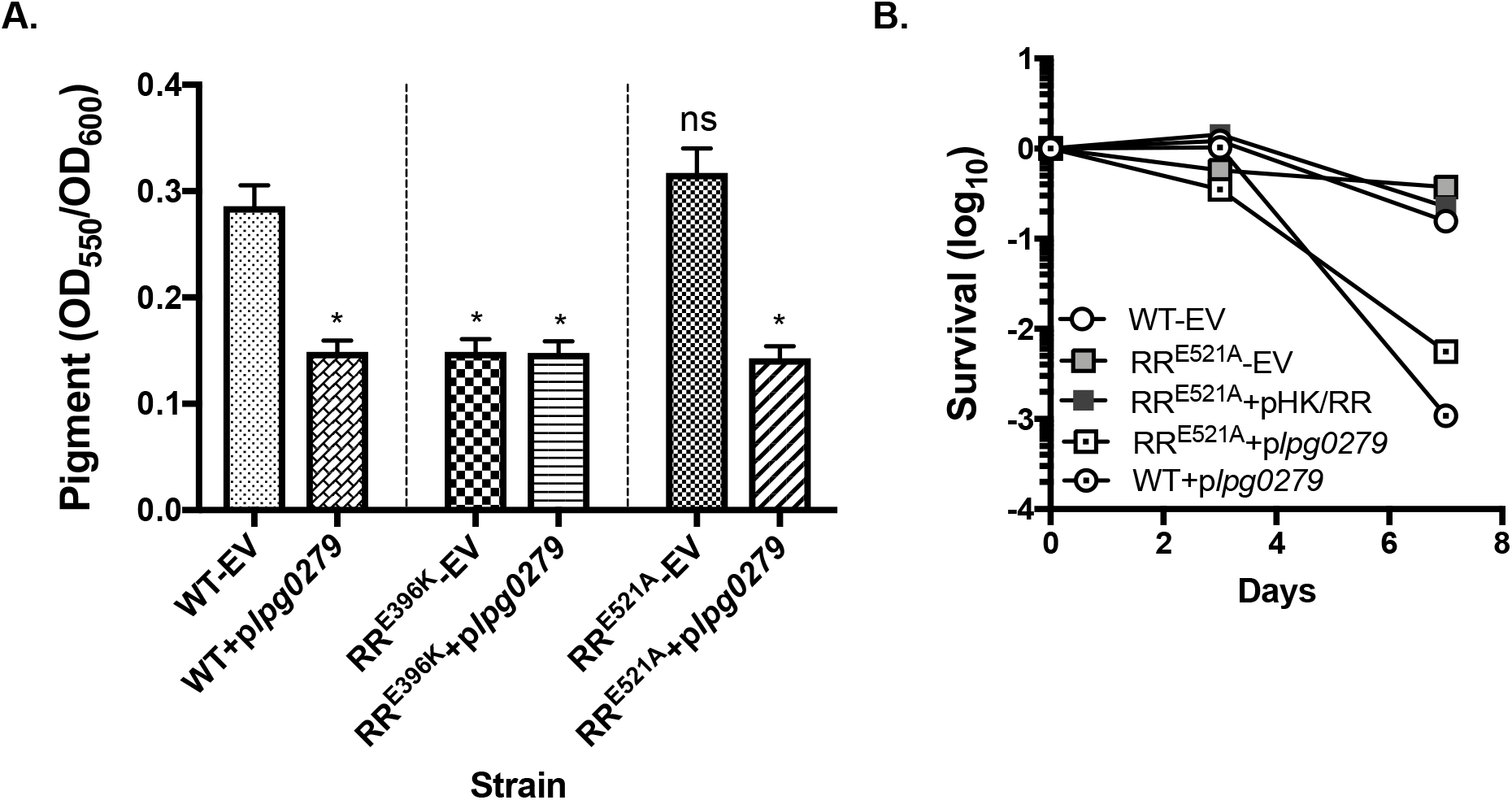
Constitutively expressed *lpg0279* is epistatic to RR^E521A^ PDE mutation. **(A)** Pigment accumulation by PE phase WT *L. pneumophila* and the Lpg0277 RR^E396K^ CDG and RR^E521A^ PDE point mutants that do or do not constitutively express p*lpg0279*. Shown are the means ± SE of four samples and are representative of results obtained in one other independent experiment. The Mann-Whitney test was used to determine statistically significant differences in pigmentation compared to WT (**ns**: not significant; **\***, *p* <0.05). **(B)** Survival of WT *L. pneumophila* and the RR^E521A^ PDE mutant that do or do not constitutively express p*lpg0279* or the complementing pHK/RR plasmid. All strains were cultured in AYE to E phase, resuspended in CDM without L-cystine, and incubated at 37°C on an orbital shaker for up to 7 days. At the times shown, culture aliquots were serially diluted for CFU enumeration on CYET. Symbols shown are the ratio of CFU(day)/CFU(titer) of duplicate samples, and data is representative of results obtained in three independent experiments. **EV:** strain harbors the pMMB206cam empty vector.

## DISCUSSION

As an intracellular pathogen, *L. pneumophila* has evolved multiple mechanisms to survive and replicate in a wide variety of environments, ranging from freshwater protozoans and human lung macrophages to nutrient-poor natural or engineered water systems. To thrive in such diverse conditions, *L. pneumophila* responds to environmental stimuli by alternating between distinct cell types. Amino acid or fatty acid starvation triggers replicating *L. pneumophila* to transition to a highly motile and infectious transmissive form, and prolonged starvation stimulates further development to the hardy MIF cell type (5, 10, 13, 47). Using an *in vitro* culture model to analyze the switch between replicative, transmissive, and resilient cell types, here we identify as a regulatory component an operon designed to regulate cyclic-di-GMP metabolism. This operon consists of *lpg0279*, which codes for a protein abundant in MIF cells (6), and *lpg0278-lpg0277*, which encodes a two-component system (TCS) (27). Together, Lpg0279 and the TCS equip *L. pneumophila* to respond to nutrient deprivation by differentiating to a non-replicative cell type that generates pigment, accumulates PHB storage granules, and maintains viability.

The *lpg0279-77* operon is induced by the stationary phase sigma factor RpoS in response to nutrient limitation (Figs. 1-4). Indeed, to survive prolonged amino acid limitation, *L. pneumophila* require not only a functional TCS (Fig. 6B) but also RpoS (48). Thus, RpoS equips *L. pneumophila* to express factors that enhance resilience in nutrient-poor environments, in part by promoting TCS-mediated production of c-di-GMP (Figs. 3 and 6) (48–51).

One factor that promotes persistence of environmental *L. pneumophila* is secretion of the pigment pyomelanin, which occurs in late PE phase. Derived from polymerization of homogentisic acid (HGA), this soluble pigment not only protects *L. pneumophila* from the damaging effects of light (42), but it also possesses ferric reductase activity that contributes to iron uptake (43). A second factor that increases environmental persistence of *L. pneumophila* is poly-3-hydroxybutyrate (PHB). To generate a reserve energy source, *L. pneumophila* increases production of PHB lipid granules at the transition to PE phase (44, 45). Formation of this energy store involves multiple enzymatic steps, and *L. pneumophila* encodes multiple PHB biosynthesis genes. The Lpg0278/Lpg0277 TCS equips *L. pneumophila* to respond to nutrient deprivation by supporting robust PHB accumulation (Fig. 6A); however, additional regulators likely contribute to the process, since *L. pneumophila* that lack the RR still generate some PHB, as judged by Nile Red fluorescence (Fig. 6). Future studies can identify the mechanism of TCS-mediated activation of PHB biosynthesis and the downstream effector of RR-generated c-di-GMP. Candidates for the TCS regulon include *L. pneumophila* genes induced in response to nutrient-limiting conditions (52).

A second messenger molecule, c-di-GMP is a wide-spread regulator of multiple bacterial physiological processes, including biofilm formation, cell cycle progression, and virulence gene expression (25, 53–55). The RR encoded by *lpg0277* is a bifunctional enzyme whose DCG and PDE domains can generate and degrade c-di-GMP production, respectively (20). When *L. pneumophila* Philadelphila-1 cells experience nutrient deprivation, activation of the TCS is predicted to increase c-di-GMP levels, based on several genetic tests of RR function. In particular, point mutations in either the RR DGC domain (Figs. 7A and C, S2B) or phosphoacceptor site (Figs. S1, S2C) phenocopy the ΔRR mutant (Figs. 5B and D, 6), whereas the PDE domain point mutant resembles WT (Figs. 7B and C, S2A). Our observations are consistent with the studies by Pecastings and colleagues of this locus in the *L. pneumophila* Lens strain: after 5 days culture on solid bacteriology medium, mutants lacking the homologous RR Lpl0329 contain less intracellular c-di-GMP than do WT cells (22). Thus, in non-replicating *L. pneumophila* cells, the DCG activity of RR Lpl0329 likely predominates. On the other hand, using proteins purified from the *L. pneumophila* Lens strain, Levet-Paulo and colleagues demonstrated that phosphorylation of the RR Lpl0329 reduced its DCG activity but left PDE activity unaltered (27). These biochemical experiments suggest that the TCS phosphorelay can decrease the local c-di-GMP level. Perhaps these differences between the *in vivo* and *in vitro* studies indicate that the enzymatic activity of RR Lpl0329 can be modulated not only through phosphorylation by its cognate HK, but also by another regulatory factor that does not co-purify with the HK or RR proteins.

One factor that does functionally interact with the TCS is Lpg0279, a protein that is conserved among *L. pneumophila*, abundant in MIF cells (6), and encoded on the *lpg0279-0277* mRNA (Fig. 1). Consistent with a function in MIF cells, *L. pneumophila* do not require Lpg0279 to transition from E to PE phase in broth. However, constitutive expression of *lpg0279* prevents replicating WT *L. pneumophila* from differentiating to the PE transmissive form, as does loss of TCS function (Fig. 5). Moreover, genetic epistasis tests predict that the MIF protein Lpg0279 acts upstream of the TCS, repressing its activity by a mechanism not yet known (Fig. 8, S3B).

One clue to Lpg0279 function is its <u>F</u>-box and <u>I</u>ntracellular <u>S</u>ignal <u>T</u>ransduction (FIST) domain, first recognized in 2007 as a component of signaling pathways in diverse prokaryotic and eukaryotic species (56). In *Pseudomonas aeruginosa*, the FIST domain of protein Pa1975 (NosP) senses nitric oxide and inhibits its co-cistronic HK to promote biofilm dispersal (57). In *L. pneumophila*, another nitric oxide sensor, the Haem-Nitric oxide/Oxygen binding protein Hnox1, is genetically and functionally linked to a GGDEF-EAL protein, Lpg1057; together this protein pair regulates biofilm formation (58). By analogy to these two regulators of bacterial differentiation, a model that warrants testing is that Lpg0279 negatively regulates TCS production of c-di-GMP in response to nitric oxide stress.

Considering our genetic data in the context of the current literature, we favor the following working model for the signal transduction pathway encoded by *lpg0279-77* (Fig. 9). When nutrients become scarce, the stationary phase sigma factor RpoS induces transcription of the *lpg0279-77* operon. As replicating *L. pneumophila* begin to transition to the PE transmissive phase, the Lpg0279 protein initially suppresses TCS production of cyclic-di-GMP. Then in response to additional stress, HK phosphorylates the RR thereby stimulating its DCG activity. An accumulation of cellular c-di-GMP promotes further progression into PE phase, production of pigment and PHB, and survival in nutrient-poor conditions.

**Fig. 9.**
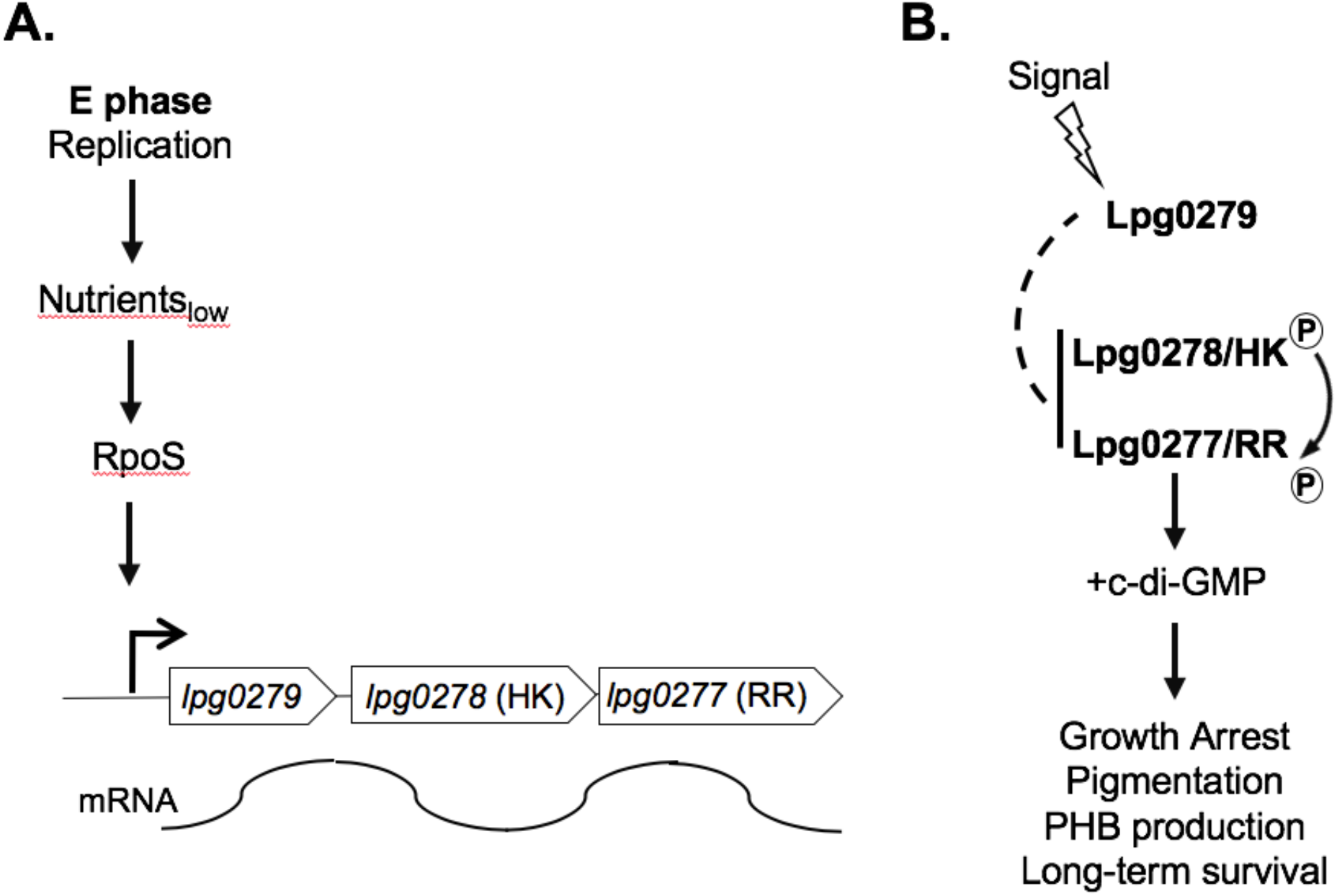
Model for *lpg0279-77* regulation of *L. pneumophila* differentiation. **(A)** When amino acids become limiting, RpoS equips replicating *L. pneumophila* to induce transcription of the *lpg0279*-*0277* operon which encodes a putative repressor and a Two Component System. **(B)** Initially Lpg0279 suppresses signaling by the Lpg0278/0277 TCS. In response to prolonged environmental stress, the TCS is derepressed and HK Lpg0278 phosphorylates RR Lpg0277. The DCG domain of RR Lpg0277 generates c-di-GMP, which arrests *L. pneumophila* replication, triggers production of pigment and PHB storage granules, and promotes survival in nutrient-poor environments.

This study extends the understanding of the regulatory circuit that governs the *L. pneumophila* life cycle. When nutrients become limiting within host cells, the stringent response alarmone ppGpp coordinates differentiation of intracellular *L. pneumophila* to a motile, infectious form equipped for transmission between host cells. The second messenger cyclic-di-GMP promotes *L. pneumophila* differentiation into a cell type equipped for persistence in nutrient poor environments. Defining c-di-GMP regulatory networks is a challenging endeavor, due to the spatial and temporal sequestering of c-di-GMP signaling, as well as the multiple enzymes that contribute to c-di-GMP metabolism in *L. pneumophila* and other bacteria (25). Accordingly, this genetic analysis of the signal transduction system comprised of the Lpg0279 MIF protein and the Lpg0278-Lpg0277 TCS can guide future molecular and biochemical studies to delineate how c-di-GMP promotes resilience of environmental *L. pneumophila*.

## MATERIALS AND METHODS

### Bacterial strains and culture conditions

The bacterial strains utilized in this study are listed in Supplementary Material Table S1. Except where indicated, all *L. pneumophila* strains were cultured in ACES (Sigma)-buffered yeast extract broth (pH = 6.9) supplemented with 0.1 mg/ml thymidine, 0.4 mg/ml L-cysteine, and 0.135 mg/ml ferric nitrate (AYET) or on solid medium containing AYET supplemented with 15 g/L agar and 2 g/L charcoal (CYET). Chemically defined medium (CDM) was prepared as previously described (38), except that ferric pyrophosphate and either L-cysteine, L-cystine, L-methionine or L-serine were omitted where indicated. Where necessary for plasmid maintenance, media were supplemented with chloramphenicol (5 µg/ml) and/or kanamycin (10 µg/mL). All *E. coli* strains were cultured using Luria-Bertani (LB) broth or agar, supplemented where necessary with ampicillin (100 ug/ml), chloramphenicol (25 µg/ml) or kanamycin (25 µg/ml). To induce gene expression from the pMMB206cam plasmid, 250μM isopropyl β-D-1-thiogalactopyranoside (IPTG; Gold Biotechnology) was added to growth media.

Bacteria from frozen stocks were struck onto CYET plates every 1-2 weeks and incubated at 37°C for ≥ 3 d until colonies developed. For experiments, colonies were inoculated into AYET and cultured overnight at 37°C on an orbital shaker to exponential (E) phase (OD_600_ < 2.5) and then subcultured in AYET for a second overnight incubation until the desired growth stage: E phase or post-exponential (PE) phase (OD_600_ > 3.5).

### Plasmids and primers

All plasmids and primers utilized in this study are listed in Supplementary Material Table S2. Plasmid p*lpg0279-gfp* was constructed by amplifying the 832 bp directly 5’ of the *lpg0279* ORF using primers EH21 and EH43, which encode *Bam*HI and *Xba*I restriction sites, respectively. After digestion, the fragment was then ligated into the GFP reporter plasmid pBH6119 5’ of a promoterless *gfpmut3* gene (13, 29). Plasmids pHK/RR and p*lpg0279* were constructed by amplifying either a 3.6 kb fragment containing *lpg0278* through *lpg0277* using primers EH69 and EH70 or a 1.2 kb fragment containing *lpg0279* using primers 79OE-F and 79OE-R, which each encode *Bam*HI and *Hind*III restriction sites; primer 79OE-F also encodes an optimal ribosome binding site (20). Following restriction enzyme digestion, the fragments were ligated into the IPTG-inducible plasmid pMMB206cam. Correct placement and orientation of the insert was verified by PCR and/or DNA sequencing.

### Mutant strain construction

The laboratory strain Lp02, a thymidine auxotroph derived from the clinical isolate Philadelphia-1 (59), was utilized as the parent strain for all constructs. Deletion mutants were generated by homologous recombination as previously described (60) using the primers listed in Table S2. The genes of interest along with ∼700 bp of 3’ and 5’ flanking DNA were amplified and cloned into the vector pGEM-T Easy (Promega) to create pGEM*lpg0277*, pGEM*lpg0280*, and pGEM*lpg0279*. The kanamycin cassette from pKD4 was amplified using primers comprised of the oligos PO and P2 along with ∼36 bp of DNA sequence homologous to the regions 3’ and 5’ of the each gene of interest. Following allelic exchange in the *E. coli* λ-red recombinase strain DY330, candidate colonies were screened by PCR and transformed into *E. coli* host strain DH5α. Point mutants RR^E396K^, RR^E521A^, and RR^D87N^ were created using the QuikChange XLII Site-Directed Mutagenesis Kit (Agilent) with plasmid pGEM*lpg0277* serving as a template and using primers sets E396K-F/E396K-R, E521A-F/E521A-R, and D87N-F/D87N-R, respectively. The recombinant alleles (*lpg0277::kan*, *lpg0278::kan* and *lpg0279::kan*) and point mutant alleles were amplified by PCR using each relevant primer pair (77del-F/77del-R, 78del-F/78del-R or 79del-F/79del-R) and introduced into Lp02 by natural transformation. Where indicated in Table S2, the kanamycin cassette was subsequently excised by Flp recombinase as previously described (61). All mutations were confirmed by DNA sequencing.

Transformation with the plasmids identified in Table S1 was conducted by electroporating isolated plasmid DNA (Qiagen) into 50 μl competent cells at 1.8 kV, 100 W and 25 μF using 1 mm cuvettes. Cells were then transferred to 950 μL AYET and incubated at 37°C for 1 h on an orbital shaker before plating on selective media. Also constructed were control strains that carry the corresponding pBH6119 or pMMB206cam empty vector.

### RNA isolation

To isolate RNA for analysis, 0.5-1.0 ml of bacterial culture at OD_600_ > 3.0 was collected by centrifugation at 12,000 × g. The pellet was resuspended in an equal volume of TRIzol reagent and then purified using the Direct-zol RNA MiniPrep kit (Zymo Research). All RNA preparations were treated with DNase I Amplification Grade or Turbo DNA-free (Invitrogen), and absence of genomic DNA was confirmed by PCR and gel electrophoresis.

### End-point PCR experiments

To determine whether *lpg0279*, *lpg0278*, and *lpg0277* are co-transcribed, 800 ng of purified RNA was used as a template to generate cDNA with the iScript cDNA Synthesis Kit (Bio-Rad). End-point PCR was then conducted using primer sets EH13/EH14 and EH1/EH2, which span the *lpg0279-lpg0278* and *lpg0278-lpg0277* intragenic regions, respectively. For the end-point PCR experiment examining co-transcription of *lpg0280* and *lpg0279*, cDNA synthesis was coupled with PCR amplification using 800 ng RNA, primer set EH55/56, and the SuperScript III One-Step RT-PCR System with Platinum *Taq* DNA Polymerase (Invitrogen). For all experiments, genomic DNA was used as a positive control, and reactions omitting the reverse transcriptase enzyme served as a negative control.

### Growth curves

Bacterial growth kinetics were analyzed by culturing *L. pneumophila* to E or PE phase as indicated, then collecting 1 ml aliquots by centrifugation at 5,000 × g for 5 minutes. The pellet was resuspended to an OD_600_ of 0.1 in 1 ml fresh AYET supplemented with chloramphenicol and ITPG, and 250 μl aliquots were dispensed into triplicate wells of a sterile 100x Honeycomb Plate (Fisher Scientific). The plates were transferred to a Bioscreen C plate reader and incubated for 36 h at 37°C with continuous shaking, with OD_600_ measurements taken at 3 h intervals.

### Pigmentation

To analyze pigment production, strains were cultured as described above to PE phase and then incubated at 37°C for an additional 1-3 days. Next, 0.5 ml samples were centrifuged at 16,000 × g for 5 min, 200 μl aliquots of each supernatant were placed in a 96-well plate, and then their absorbance at OD_550_ was quantified on a plate reader. To normalize pigment values to cell density, each cell pellet was resuspended in PBS to its original volume, and then the OD_600_ of 100 μl aliquots was quantified on a plate reader. All measurements were performed in duplicate.

### GFP transcriptional reporter experiments

To analyze activity of the *lpg0279-0277* promoter, strains EH224, EH97 and EH102 which each harbor plasmid p*lpg0279-gfp* were cultured overnight to E phase, and then diluted to an OD_600_ of 0.4-0.8 in either AYET or CDM that lacked L-cysteine, L-serine or L-methionine, as indicated. The AYET and CDM bacterial suspensions were supplemented aseptically with 0.135 mg/ml ferric nitrate and/or 2.27 mM, 1.14 mM or 0.7 mM L-cysteine, as indicated. All cultures were then further incubated at 37°C for 10-12 h on an orbital shaker. Measurements were taken at 2-3 h intervals by centrifuging 800 μl aliquots, resuspending the pellet in an equal volume of PBS, and quantifying fluorescence of triplicate 200 μl samples at 485_EX_/528_EM_ on a Biotek plate reader. To normalize all fluorescence readings to cell density, the OD_600_ of a 1/10 dilution of each cell suspension was quantified with a spectrophotometer.

### PHB measurement by Nile Red staining

To analyze intracellular lipid (PHB) content, 4-6 ml aliquots of E phase cultures were first collected by centrifugation (5 min at 5,000 × g) and the cell pellets resuspended in an equal volume of CDM supplemented with thymidine (0.1 mg/ml), chloramphenicol (25 µg/ml), and IPTG (250 μM), but lacking L-cysteine and L-cystine. Cultures were then incubated for 24 h at 37°C on an orbital shaker. PHB content was quantified for the initial E phase cultures and again following the 24 h incubation using the fluorescent dye Nile Red (Invitrogen) as described (45), with the following modifications. Briefly, aliquots of bacterial cultures were collected by centrifugation and resuspended in an equal volume of deionized water before fixing the cells with 1% (v/v) formaldehyde at room temperature for 30 min. After washing to remove the formaldehyde, cell density was adjusted to OD_600_ 0.5 in 1 ml of deionized water, and the cells were stained by adding 1 μl of a 25 mM Nile Red stock solution suspended in DMSO. The cells were incubated at room temperature in the dark for 1 h, and then 200 μl aliquots were measured in triplicate on a Biotek plate reader at 545_EX_/600_EM_.

### Survival assay

To assess long-term survival of *L. pneumophila* in the absence of L-cysteine, cultures prepared as for PHB measurement described above were incubated at 37°C on an orbital shaker for 7 days. At the times indicated, duplicate samples were removed, serially diluted, and plated to enumerate CFUs on CYET.

## ACKNOWLEDGEMENTS

This work was supported by the Michigan Predoctoral Genetics Training Program (NIH T32-GM-007544; E.D.H.), a University of Michigan Rackham Graduate Student Research Grant (E.D.H.), and the Endowment for Basic Sciences at the University of Michigan Medical School (M.S.S.).

## SUPPLEMENTAL TABLES

**Table S1.** Strains used in this study.

| Strain | Relevant properties | Source |
| --- | --- | --- |
| ***E. coli*** |  |  |
| DH5α | *supE44 ΔlacU169 (80 lacZΔM15) hsdR17 recA1 endA1 gyrA96 thi-1 relA1* | Laboratory collection |
| DY330 | W3110 *ΔlacU169 gal490 λcI857 Δ(cro-bioA)* | (62) |
| EH207 | DH5α *plpg0279-gfp* | This work |
| EH272 | DH5α *plpg0279-77* | This work |
| EH140 | DH5α *plpg0279* | This work |
| ***L. pneumophila*** |  |  |
| MB110 | Lp02 wild type; *thyA hsdR rpsL* (Str<sup>r</sup>) | (63) |
| EH284 | Lp02 pMMB206cam | This work |
| EH286 | Lp02 *Δlpg0277::FRT* pMMB206cam | This work |
| EH276 | Lp02 *Δlpg0277::FRT plpg0278-77* | This work |
| EH357 | Lp02 *Δlpg0278::FRT* pMMB206cam | This work |
| EH352 | Lp02 *Δlpg0278::FRT plpg0278-77* | This work |
| EH350 | Lp02 *Δlpg0279::FRT-kan-FRT* pMMB206cam | This work |
| EH160 | Lp02 *Δlpg0279::FRT-kan-FRT plpg0279* | This work |
| EH151 | Lp02 *plpg0279* | This work |
| EH344 | Lp02 Lpg0277<sup>E396K</sup> pMMB206cam | This work |
| EH311 | Lp02 Lpg0277<sup>E396K</sup> *plpg0278-77* | This work |
| EH346 | Lp02 Lpg0277<sup>E521A</sup> pMMB206cam | This work |
| EH314 | Lp02 Lpg0277<sup>E521A</sup> *plpg0278-77* | This work |
| EH102 | Lp02 *pflaA-gfp* | This work |
| EH224 | Lp02 *plpg0279-gfp* | This work |
| EH97 | Lp02 pBH6119 | This work |
| MB410 | Lp02 *ΔfliA::kan* | (14) |
| MB443 | Lp02 *ΔrpoS::kan* | (34) |
| MB696 | Lp02 *ΔrelA::kan* | (64) |
| MB413 | Lp02 *ΔletA::kan* | (14) |
| MB416 | Lp02 *ΔletS::kan* | (14) |
| EH358 | Lp02 *ΔfliA plpg0279-gfp* | This work |
| EH360 | Lp02 *ΔrpoS plpg0279-gfp* | This work |
| EH366 | Lp02 *ΔrelA plpg0279-gfp* | This work |
| EH362 | Lp02 *ΔletA plpg0279-gfp* | This work |
| EH364 | Lp02 *ΔletS plpg0279-gfp* | This work |
| MB1230 | Lp02 *ΔletA::kan pflaA-gfp* | (14) |
| EH370 | Lp02 *ΔletS::kan pflaA-gfp* | This work |
| MB1234 | Lp02 *ΔrpoS::kan pflaA-gfp* | (34) |
| MB1014 | Lp02 *ΔrelA::kan pflaA-gfp* | (64) |
| EH368 | Lp02 *ΔfliA::kan pflaA-gfp* | This work |
| EH373 | Lp02 Lpg0277<sup>E521A</sup> *plpg0279* | This work |
| EH375 | Lp02 Lpg0277<sup>E396K</sup> *plpg0279* | This work |

**Table S2.**
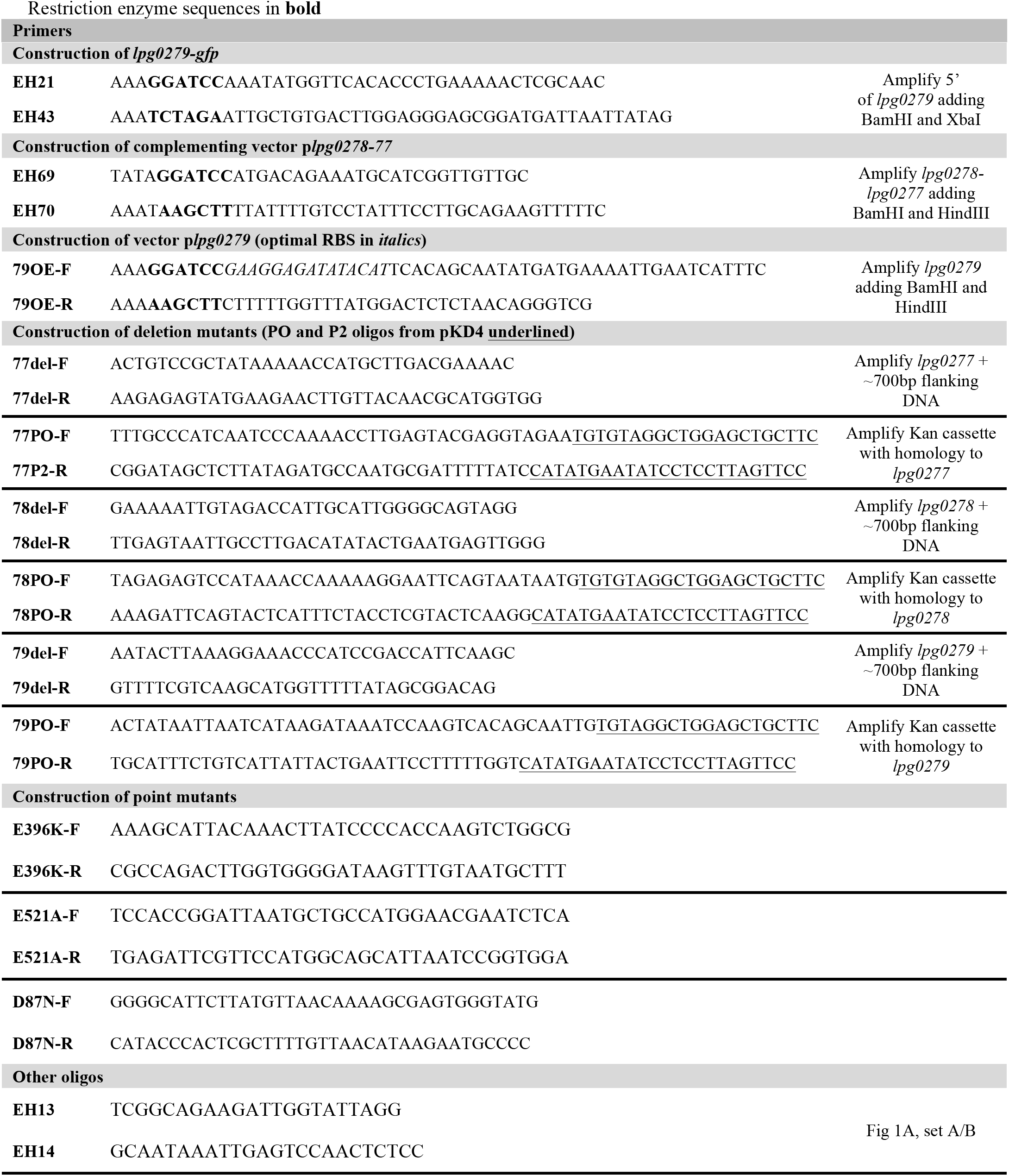
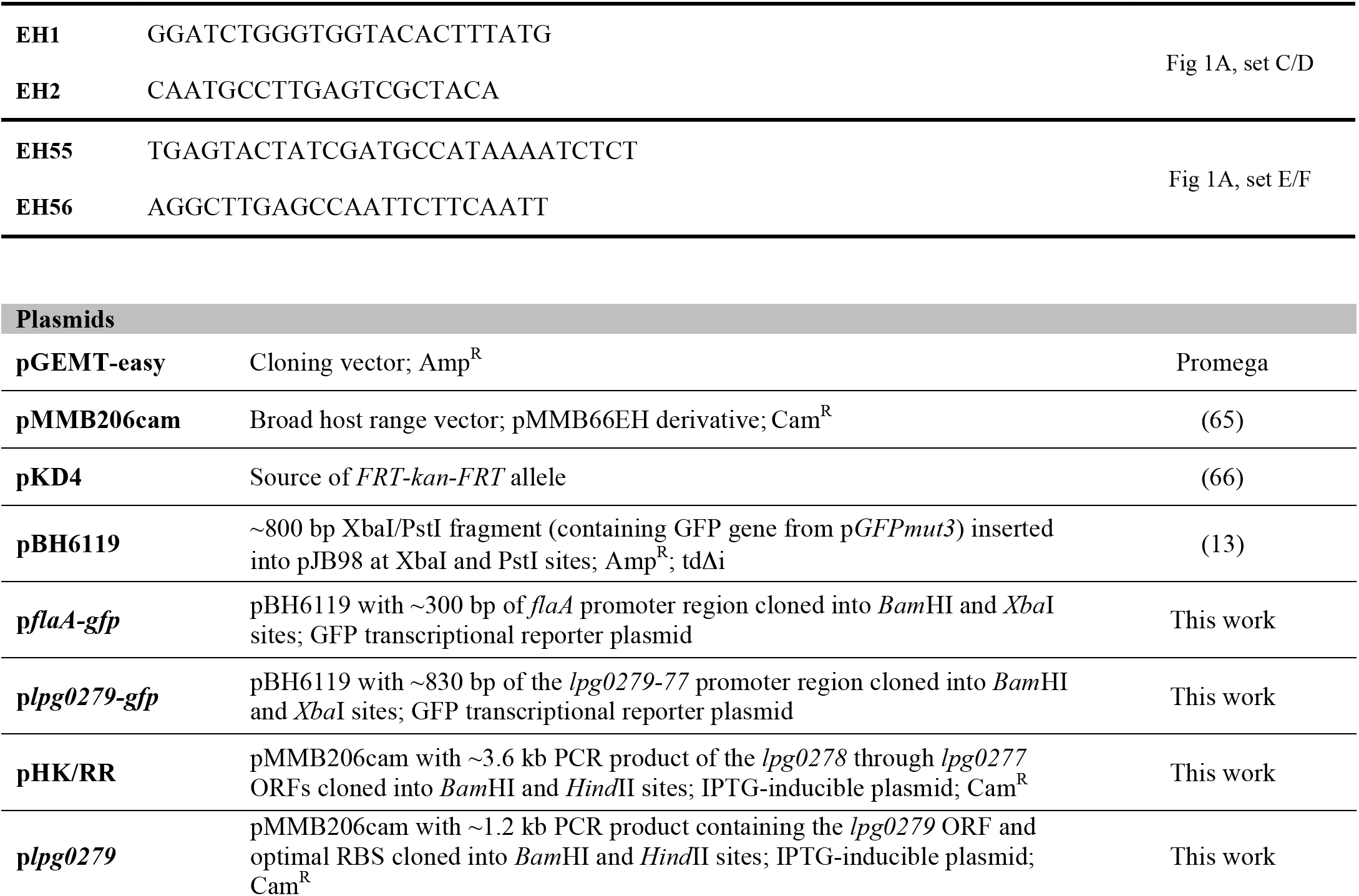
Primers and plasmids used in this study.

## SUPPLEMENTAL FIGURES

**Fig S1.**
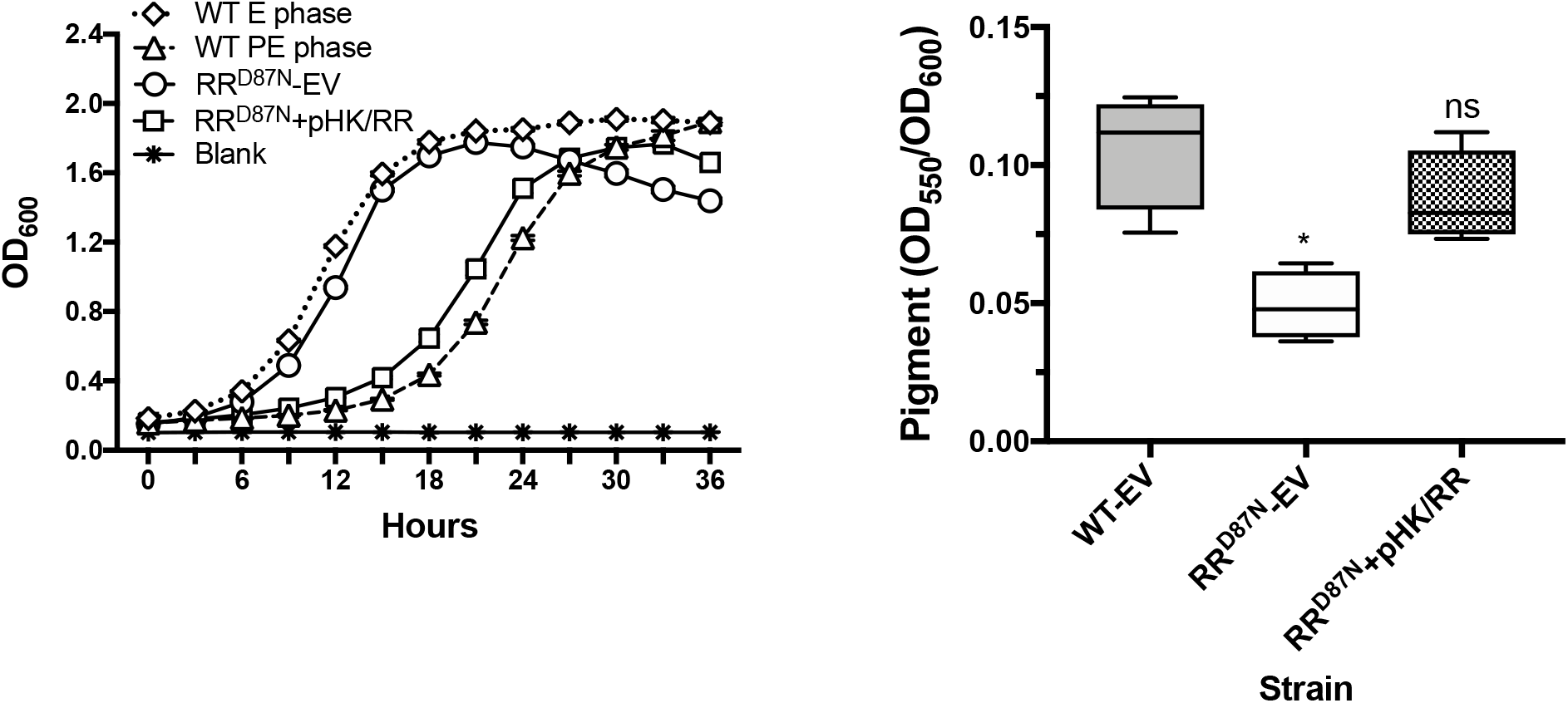
The Lpg0277 RR phosphoacceptor site promotes *L. pneumophila* differentiation to the PE phase. Growth kinetics and pigment production by WT *L. pneumophila* and RR^D87N^ phosphoacceptor site point mutant and the corresponding complemented strain. Results shown are representative of one other independent experiment. The Mann-Whitney test was used to determine statistically significant differences in pigmentation compared to WT (**ns**: not significant; **\***, *p* <0.05). **EV:** strain harbors the pMMB206cam empty vector.

**Fig. S2.**
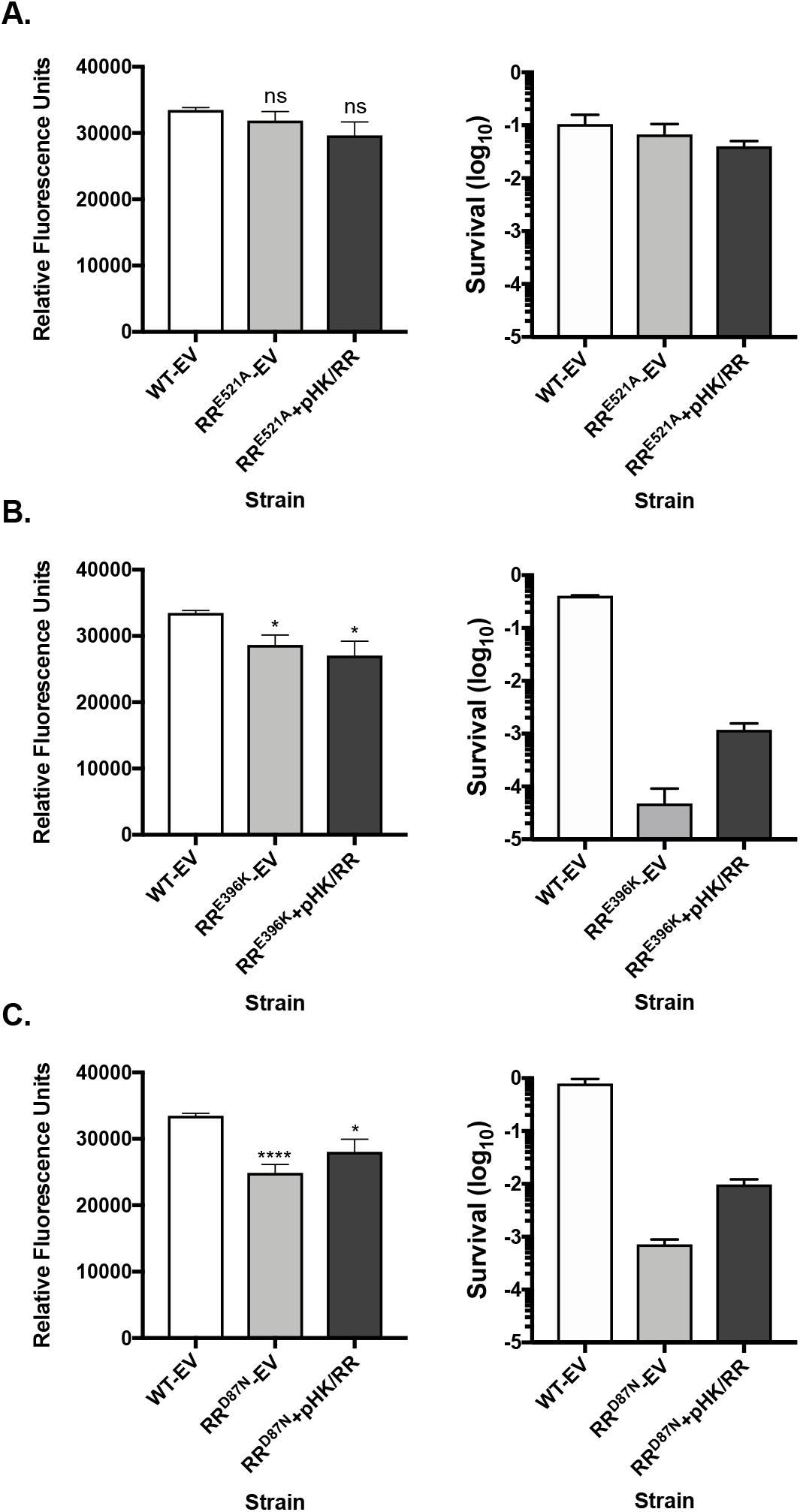
The GGDEF domain and phosphoacceptor site of the RR contribute to PHB production and promote survival during prolonged nutrient deprivation. PHB quantification and survival of E phase cultures after 24 h and 7 d L-cysteine deprivation, respectively. WT *L. pneumophila* were compared to (A) RR^E521A^ PDE, (B) RR^E396K^ DCG, and (C) RR^D87N^ phosphoacceptor site point mutants, and the respective complemented strains bearing plasmid pHK/RR. PHB values represent the means ± SE of pooled Nile Red fluorescence data from triplicate samples in four independent experiments. A two-tailed Student’s t-test was used to determine statistically significant differences in fluorescence compared to WT-EV (**ns**: no significance; **\***, *p* < 0.05; **\*\*\*\***, *p* < 0.0001). Survival data is the ratio of CFU(day 7)/CFU(titer), and data is representative of results obtained in two or more independent experiments. **EV**: strain harbors the pMMB206cam empty vector.

**Fig. S3.**
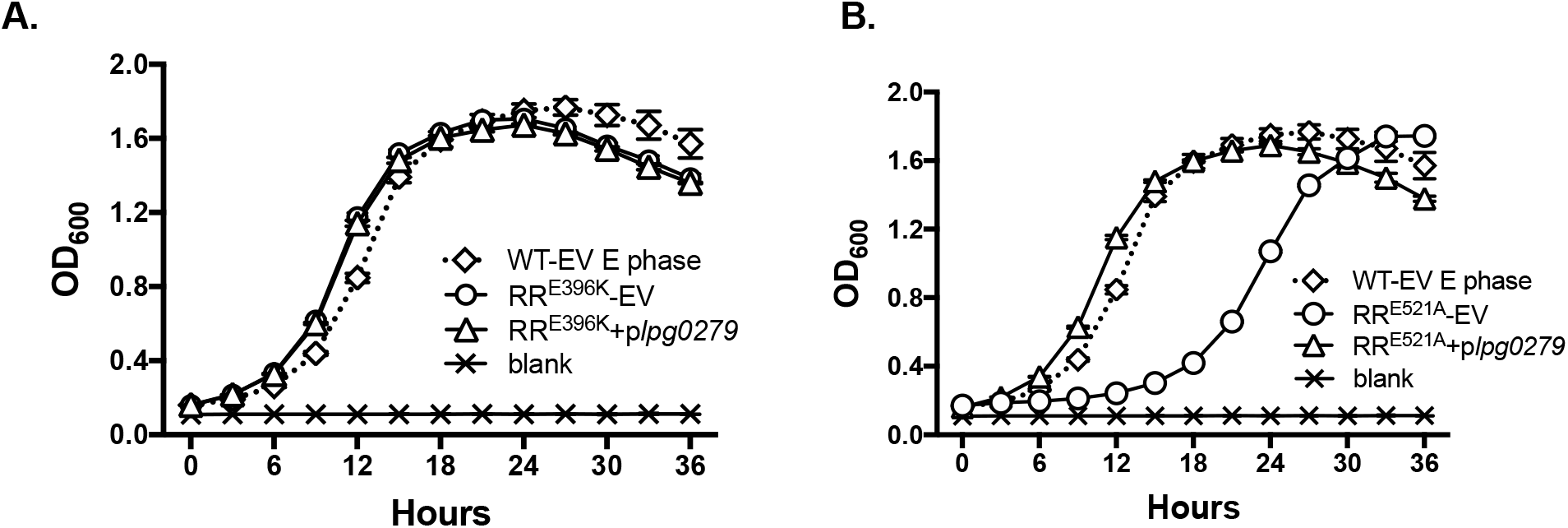
Disruption of the Lpg0277 RR^E521A^ PDE domain is not sufficient to suppress the differentiation defect of PE phase *L. pneumophila* constitutively expressing *lpg0279*. Growth kinetics of PE phase WT *L. pneumophila* and the **(A)** Lpg0277 RR^E396K^ DCG and **(B)** RR^E521A^ PDE point mutants that harbor either p*lpg0279* or pMMB206 empty vector (**EV**). Symbols denote the means ± SE of triplicate samples, and data shown are representative of one additional independent experiment.

